# Native structure of the RhopH complex, a key determinant of malaria parasite nutrient acquisition

**DOI:** 10.1101/2021.01.10.425752

**Authors:** Chi-Min Ho, Jonathan Jih, Mason Lai, Xiaorun Li, Daniel E. Goldberg, Josh R. Beck, Z. Hong Zhou

**Author notes:** Correspondence and request for materials should be addressed to CMH.

## Abstract

The RhopH complex is implicated in malaria parasites’ ability to invade and create new permeability pathways in host erythrocytes, but its mechanisms remain poorly understood. Here we enrich the endogenous RhopH complex in a native soluble form, comprising RhopH2, CLAG3.1 and RhopH3, directly from parasite cell lysates and determine its atomic structure using cryo electron microscopy, mass spectrometry, and the cryoID program. This first direct observation of an exported *P. falciparum* transmembrane protein—in a soluble, trafficking state and with atomic details of buried putative membrane-insertion helices—offers insights into assembly and trafficking of RhopH and other parasite-derived complexes to the erythrocyte membrane. Our study demonstrates the potential endogenous structural proteomics approach holds for elucidating the molecular mechanisms of hard-to-isolate complexes in their native, functional forms.

## Introduction

Nearly half the world’s population is at risk of contracting Malaria. To keep the disease at bay, we rely heavily on artemisinin-based therapies, limiting Malaria’s impact to 200 million cases and half a million deaths each year^1^. The recently discovered *de novo* emergence of artemisinin-resistant malaria parasites in Africa heightens the urgent need to define, and thus target, molecular machineries essential for parasites’ survival in human erythrocytes, thereby facilitating the development of new therapeutics with novel modes of action^1–3^. To survive inside human red blood cells, the malaria parasite *Plasmodium falciparum* deploys hundreds of effector proteins that extensively remodel the normally quiescent host erythrocyte, building new infrastructure for nutrient uptake, waste efflux, and immune evasion to support the active growth and replication of the parasite^4,5^. Members of the high-molecular mass rhoptry protein complex known as the RhopH complex—comprising CLAG3.1, RhopH2 and RhopH3—have been shown to play a key role in this process by directly or indirectly contributing to the activity of the *Plasmodium* surface anion channel (PSAC), a novel ion channel that appears at the cell surface in malaria parasite-infected erythrocytes and has been shown to mediate increased permeability of the erythrocyte plasma membrane to various solutes^6–18^.

The mechanism underlying the role of the RhopH complex in PSAC activity remains poorly understood. How the components of the RhopH complex are trafficked from the parasite out to the erythrocyte membrane is also not known, although it has been hypothesized that they may assume a soluble form or associate with additional proteins that protect them as they traverse the host cell cytosol^9^. Another theory suggests that they may cross the cytosol via a series of membranous compartments created by the parasite, known as the exomembrane system^19,20^. High-resolution structures of the RhopH complex in either membrane-bound or soluble forms would help to answer these open questions and provide a much-needed structural framework to rationalize the large body of seemingly contradictory phenotypic and biochemical data that surround this intriguing complex.

To address the question of how members of the RhopH complex are delivered to the erythrocyte membrane, we sought to investigate whether a soluble complex containing one or more components of the RhopH complex exists in soluble parasite lysate, and determine its structure if it did. Unfortunately, the extreme difficulty of producing properly folded and assembled *P. falciparum* protein complexes via reconstitution or co-expression in heterologous systems has thus far stymied efforts to obtain high resolution structures of the RhopH complex. Furthermore, the uncertainty about the exact composition of a putative soluble RhopH complex precluded the use of a heterologous system to recapitulate the complex. To address challenges like this, we recently established an endogenous structural proteomics approach that combines the use of cryoEM, mass spectrometry, and cryoID, a program we developed to identify proteins in sub-4.0Å cryoEM density maps, to identify and determine the structures of unknown protein complexes (Fig. 1) ^21^. Here, we have used this approach to obtain a near-atomic resolution structure of the unmodified RhopH complex in a soluble state, enriched directly from *P.falciparum* parasite lysate. The structure explains the body of existing biochemical data, provides insights into the dual roles of the RhopH complex in both invasion and creation of new permeability pathways in host erythrocytes, and sheds light on the long-standing question of how parasite transmembrane proteins are trafficked to the erythrocyte membrane. The study also demonstrates the power of the endogenous structural proteomics approach to provide exciting new insights into the molecular mechanisms of hard-to-isolate complexes from challenging native sources where conventional structural biology approaches have failed.

**Figure 1.**
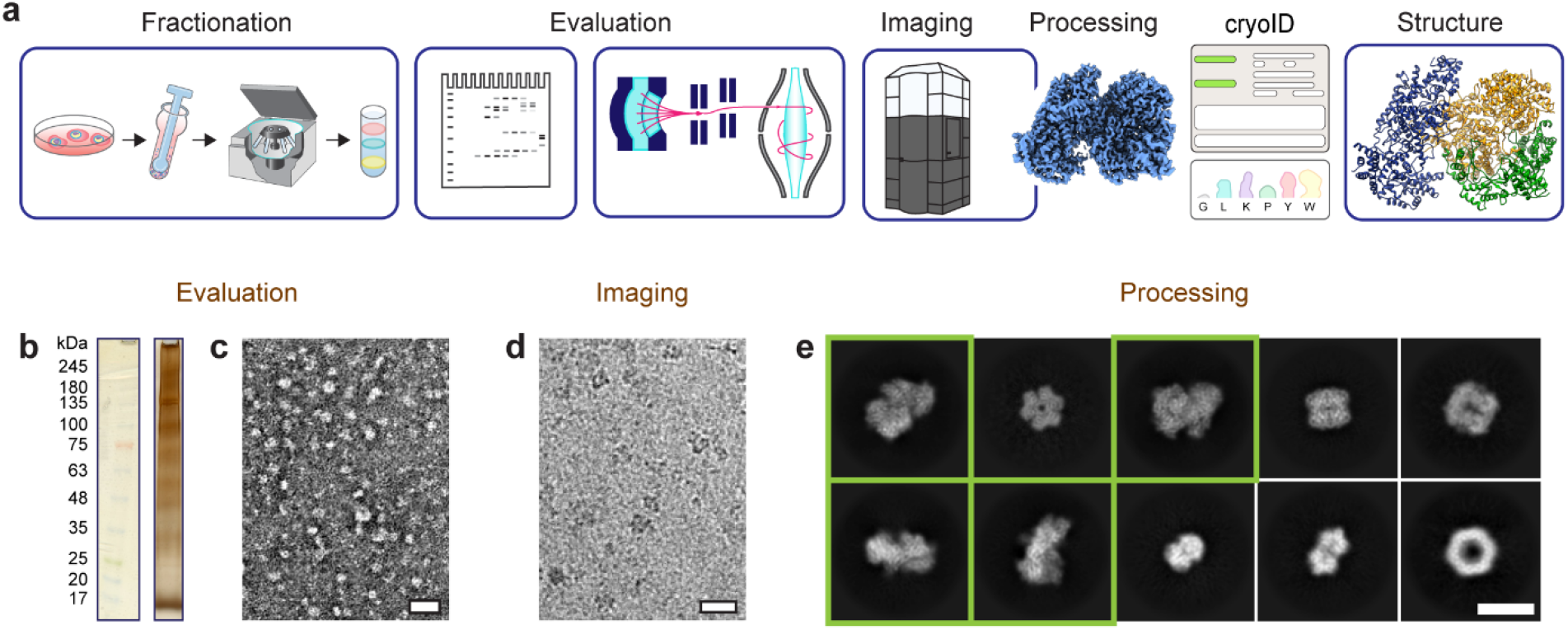
Endogenous structural proteomics workflow using cryoID. **a,** Depiction of the workflow. Plasmodium falciparum parasites and parasitophorous vacuoles were saponin-released from P. falciparum-infected red blood cells, and the resulting parasite and vacuole pellets were lysed and fractionated using a sucrose gradient. The fractions were evaluated by SDS-PAGE, mass spectrometry, and negative stain electron microscopy. CryoEM imaging and analysis of the selected fraction yielded a 3.7Å resolution cryoEM density map. The proteins in the cryoEM map were identified using cryoID and then modeled de novo, yielding the final atomic resolution structure of the RhopH complex. **b,** Silver-stained SDS–PAGE of the selected fraction of the lysate. **c-d,** Representative negative stain (**c**) and cryoEM (**d**) micrographs of the selected fraction. Scale bars, 20 nm. **e,** Representative two-dimensional class averages corresponding to multiple protein complexes present in the single dataset of cryoEM micrographs. Class averages that ultimately gave rise to the structure of the RhopH complex presented here are boxed in green. Scale bar, 10 nm.

## Results

### Identification and structure determination of soluble RhopH complex

Following an endogenous structural proteomics workflow^21^ (Fig. 1a), we used mass spectrometry to identify fractions of *Plasmodium falciparum* parasite lysate that both looked promising by negative stain EM and contained one or more components of the RhopH complex (Fig. 1b-c, Supplementary Table 1). Analysis of a cryoEM dataset collected from one of these fractions in RELION^22^ and cryoSPARC^23^ yielded numerous promising 2D class averages (Fig. 1d-e). We identified a subset of these class averages that gave rise to a near-atomic resolution cryoEM density map of a novel, asymmetric protein complex at an overall resolution of 3.7Å after *ab initio* reconstruction and non-uniform refinement in cryoSPARC (Fig. 2).

**Figure 2.**
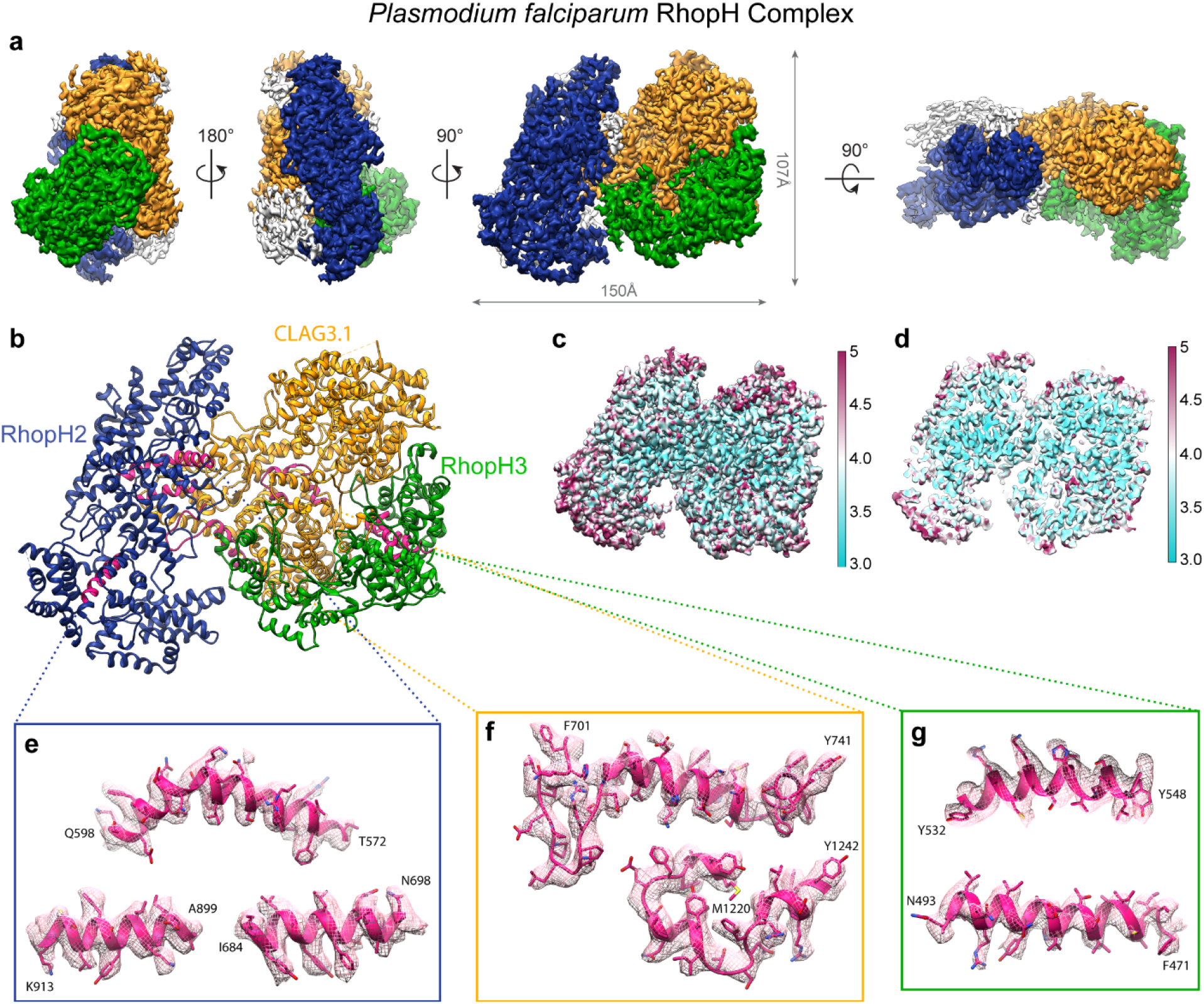
CryoEM density map of the soluble RhopH complex. **a,** CryoEM density map of the RhopH complex viewed from multiple angles, with RhopH2, CLAG3.1, and RhopH3 colored in navy, gold, and green respectively. Unmodeled regions are shown in grey. **b,** Atomic model of the soluble RhopH complex, colored as in (**a**), with the query segments used for protein identification by cryoID colored in pink. **c-d,** Local resolution evaluation of the soluble RhopH complex map, calculated using Resmap^24^, colored according to resolution, and displayed as a full surface (**c**) and central slice (**d**). **e-g,** Detailed view of each set of query sequences used for cryoID identification for RhopH2 (**e**), CLAG3.1 (**f**), and RhopH3 (**g**), shown with corresponding cryoEM density (mesh).

Following further focused classification and refinement in RELION to improve local resolutions throughout the map, we were able to use *cryoID* to successfully identify all three components of the complex as RhopH2, CLAG3.1, and RhopH3, the three members of the RhopH complex (Fig. 2b, e-g Supplementary Videos 1–3). Clear sidechain densities throughout most of the map enabled us to build *de novo* atomic models of the three components (Fig. 2b), revealing a tripartite protein complex containing a single copy of each of the three proteins, with a total calculated mass of 435kDa, assuming no post-translational truncations. Although the CLAG3.1 and CLAG3.2 isoforms share 96% protein sequence identity, there is a known hypervariable region (HVR) (F1094-G1142), where CLAG3.1 in NF54 parasites has an extra two amino acids (T1140-H1141) that are not found in CLAG3.2 in NF54 parasites. Fortunately, we had strong density in this region that enabled us to definitively resolve that the isoform in our map is CLAG3.1 (Supplementary Fig. 1). The overall map quality was also such that we were able to model the bulk of the complex’s visible density, the main exception being a small auxiliary domain near the C-terminus of RhopH2, which at a lower density threshold value (σ = 0.0355) accounts for approximately 6.9% of our complex’s total volume (Supplementary Fig. 2). Though the density here was of insufficient quality to register residues, we were able to discern that the region mainly comprises several twisted β-strands constituting a β-sheet-rich core.

CLAG3.1 forms the core of the complex and is situated between RhopH2 and RhopH3 (Fig. 2a-b, Supplementary Video 4). RhopH2 and RhopH3 both share substantial binding interfaces with CLAG3.1, but make minimal contacts with each other (Supplementary Video 4). Buried and exposed surface areas were calculated using PISA^25^. Of the two, the CLAG3.1-RhopH3 interface is more extensive, accounting for a buried surface area of 5416 Å^2^ (~11-15% of the total exposed surface area of either CLAG3.1 or RhopH3), compared with a buried surface area of 3410 Å^2^ for the CLAG3.1-RhopH2 interface (~7-7.5% of the total exposed surface area of either CLAG3.1 or RhopH2). All three protein components contain an unusually high abundance of cysteines in their primary sequence (13%, 10.5%, and 15.6% in RhopH2, CLAG3.1, and RhopH3, respectively), which is particularly striking given the relatively low presence of cysteines in the *P. falciparum* proteome (~1.7%). Our CLAG3.1 structure contains three sets of disulfide bonds, while RhopH2 and RhopH3 contain five sets of disulfide bonds each (Supplementary Fig. 3).

### RhopH3 architecture explains RhopH3 truncation phenotypes

RhopH3 is a 897 amino acid (a.a.) protein, whose visible structure can be divided into three domains (Fig. 3): an elongated N-terminal domain arising from RhopH3 exons 1-3 (corresponding to a.a. 1-378), composed of a set of interleaved β-sheets flanked by α-helices; a globular α-helix-rich “middle domain” from RhopH3 exons 4-6 (a.a. 379-675); and an extended C-terminal domain arising from RhopH3 exon 7, characterized by a long loop-rich segment that extends towards RhopH2 (Fig. 3c). The first N-terminal 25 residues were not modeled due to disordered density, nor were residues after a.a. 738, although scattered weak “pigtail” density can be observed after a.a. 738 further extending into a cleft in the RhopH complex between the major bodies of RhopH2 and RhopH3 (Supplementary Fig. 2e). The middle domain, formed by exons 4-6, is quite distant from the RhopH2-CLAG3 interface (Fig. 3a,c), explaining previous studies suggesting that RhopH3 exons 4-6 are required for trafficking to the rhoptries, while exons 1-3 and/or exon 7 are involved in both binding to CLAG3 as well as potentially helping to form the interaction between CLAG3 and RhopH2 [9, 12].

**Figure 3.**
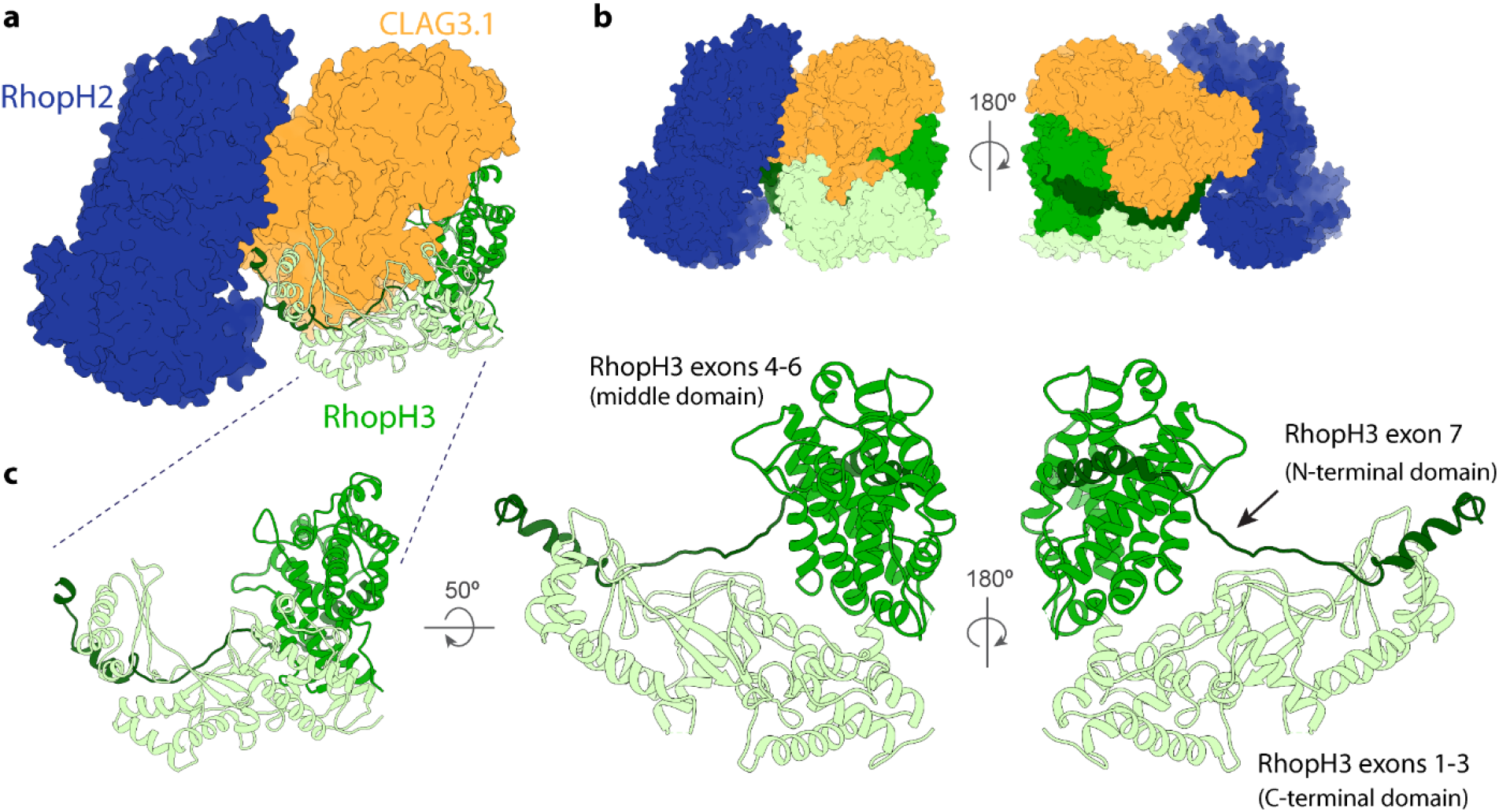
Structural details of RhopH3. **a,** RhopH3 (ribbon diagram) and RhopH2 and CLAG3.1 (space-filling surface diagram in different blue and orange, respectively) showing the interfaces between RhopH3 and the other components of the RhopH complex. **b,** Front and back views showing the arrangement of RhopH3 domains. **c,** Multiple views of the RhopH3 N-terminal, middle, and C-terminal domains, corresponding to RhopH3 exons 1-3, 4-6, and 7, and colored mint, green, and forest, respectively.

The aforementioned abundance of cysteines in the RhopH complex plays an especially important role in the structural organization of RhopH3. Notwithstanding the unmodeled auxiliary domain of RhopH2, RhopH3 is unique amongst the constituents of the RhopH complex in possessing the only β-sheets within the main complex; in contrast, RhopH2 and CLAG3.1 are dominated by α-helices. Three sets of β-sheets present in RhopH3’s N-terminal domain give rise to an interleaved β-sheet motif, largely facilitated by a 20 residue-long continuous loop (a.a. 225-245) that contributes a single β-strand to all three sets of β-sheets, effectively chaining the three β-sheets together (Supplementary Fig. 4). Notably, this 20 residue continuous loop contains two cysteine residues, Cys231 and Cys244, each of which participates in a disulfide bond [with Cys157 (from an adjacent loop that also contributes β-strands to the interleaved β-sheets) and Cys253 of RhopH3, respectively] in a manner that could fix this structurally critical β-strand-containing loop in place (Supplementary Figs. 3,4). We therefore speculate that these cysteine residues are of vital importance to the proper folding and assembly of RhopH3 (See Discussion).

### Putative transmembrane and extracellular elements of CLAG3.1 are buried

The 1209 residues of CLAG3.1 in our structure can also be divided into three domains (Fig. 4a). Residues 52-675 form a globular N-terminal domain that shares an interface with the RhopH3 “middle domain” (Fig. 4). Residues 688-978 form a squid-shaped “middle” domain, with the head sharing an interface with RhopH2, while the legs wrap around the third CLAG3.1 domain, a helical bundle comprising residues 979-1289 (Fig. 4a).

**Figure 4.**
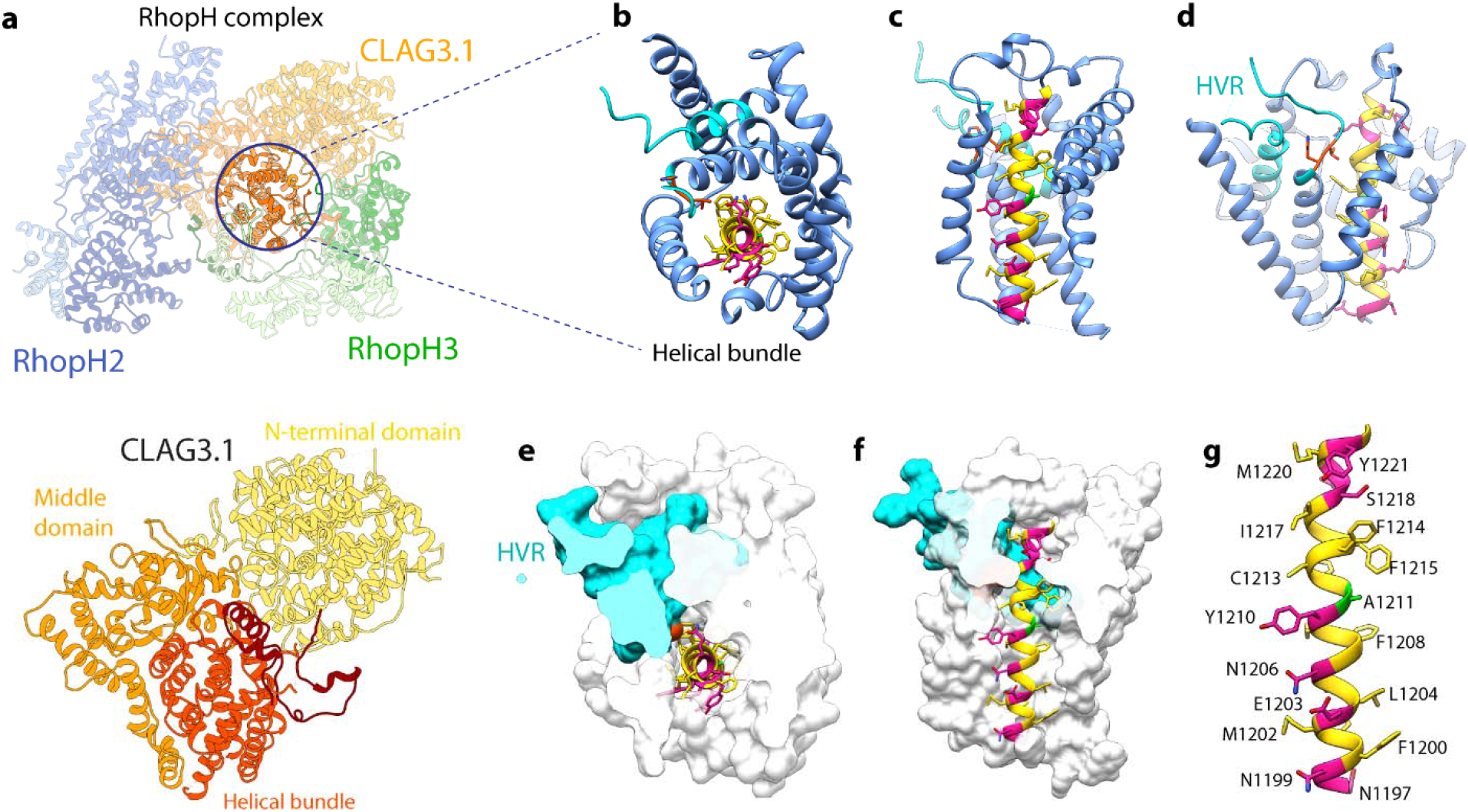
Structural details of CLAG3.1. **a,** Atomic model of CLAG3.1 shown as ribbon. **b-c,** Detailed views of the CLAG3.1 helical bundle and hypervariable region (HVR) from the top (**b**) and side (**c**). One helix in front is removed for clarity in (**c**). The helical bundle is shown in cornflower blue, the HVR in cyan. For CLAG3.1 residues N1197-A1222, hydrophobic residues, charged and polar residues, and A1211 are shown in yellow, pink, and green, respectively. The CLAG3.1-specific T1140-H1141 residues are shown in orange. **d,** View rotated 90° from (**c**), better showing the HVR. **e-f,** Cutaway space filling top (**e**) and side (**f**) views of the helical bundle and HVR, except residues N1197-A1222, which are displayed in ribbon diagram. **g,** Detailed view of CLAG3.1 N1197-A1222.

CLAG3 has been demonstrated to modulate PSAC activity^10,26^, and it has been postulated that CLAG3 may be a major or sole component of the PSAC ion-conducting pore^10^. Based on membrane protein prediction software and helical wheel analysis, CLAG3.2 residues 1203-1223 were hypothesized to constitute a transmembrane domain, with F1200-S1217 predicted to form an amphipathic helix that oligomerizes with the same helix from multiple CLAG3.2 monomers to form the transmembrane pore of PSAC, similar to oligomeric pore-forming bacterial toxins^11,27^. However, recent CLAG3.1/3.2 knockdown and complete knockout studies suggest that CLAG3 isoforms cannot be the sole component of the PSAC transmembrane pore^8,28^.

In our structure, CLAG3.1 residues 1210-1223, previously predicted to constitute a transmembrane helix, are embedded in the middle of the helical bundle formed by residues 979-1289, near the C-terminus of CLAG3 (Fig. 4b-g). As such, they are buried in the core of the RhopH complex and mostly shielded from the solvent by the helical bundle (Fig. 4b-g). Inspection of the residues corresponding to the putative transmembrane helix (a.a. 1200-1223) in the structure reveals that F1200-S1217 does form an α helix, but it is not amphipathic across the entire length of that sequence as was previously predicted (Fig. 4g)^11^. The amphipathic portion of the helix (N1199-Y1210) spans 3 turns (11 residues) and measures ~15Å in length (Fig. 4g), 4-7 residues shorter than the typical minimum length of a transmembrane helix. As typical biological membranes range from 20-40 nm in thickness, it is possible that the predicted transmembrane helix either fails to reach all the way across the membrane, or may have to form a coiled-coil interaction with another helix in order to do so. The polar face of the N-terminal 3 turns of this helix, comprising residues N1199, E1203, N1206, and possibly Y1210, are solvent exposed and contribute to the outward, solvent-facing surface of the helical bundle (Supplementary Video 5,6). All other residues in the N1199-A1222 helix are buried.

Residues Y1093-G1142 constitute the HVR previously shown to be exposed on the surface of infected red blood cells in membrane-associated CLAG3.1/3.2 ^10^ (Supplementary Fig. 1). In our structure, this region forms the ends of two helices in the helical bundle, as well as the long loop between them, which extends up from the middle of the CLAG3.1 C-terminal helical bundle through the CLAG3.1-RhopH3 N-terminal domain interface (Fig. 4 and Fig. 5a). There are 20 residues in the middle of the loop that are unstructured in our density map. Intriguingly, most of the HVR that we can see in our structure is buried deep in the middle of the helical bundle, shielded from solvent by both the CLAG3.1 helical bundle and the RhopH3 N-terminal domain (Fig. 4b-g, Fig. 5a, Supplementary Video 5,6).

**Figure 5.**
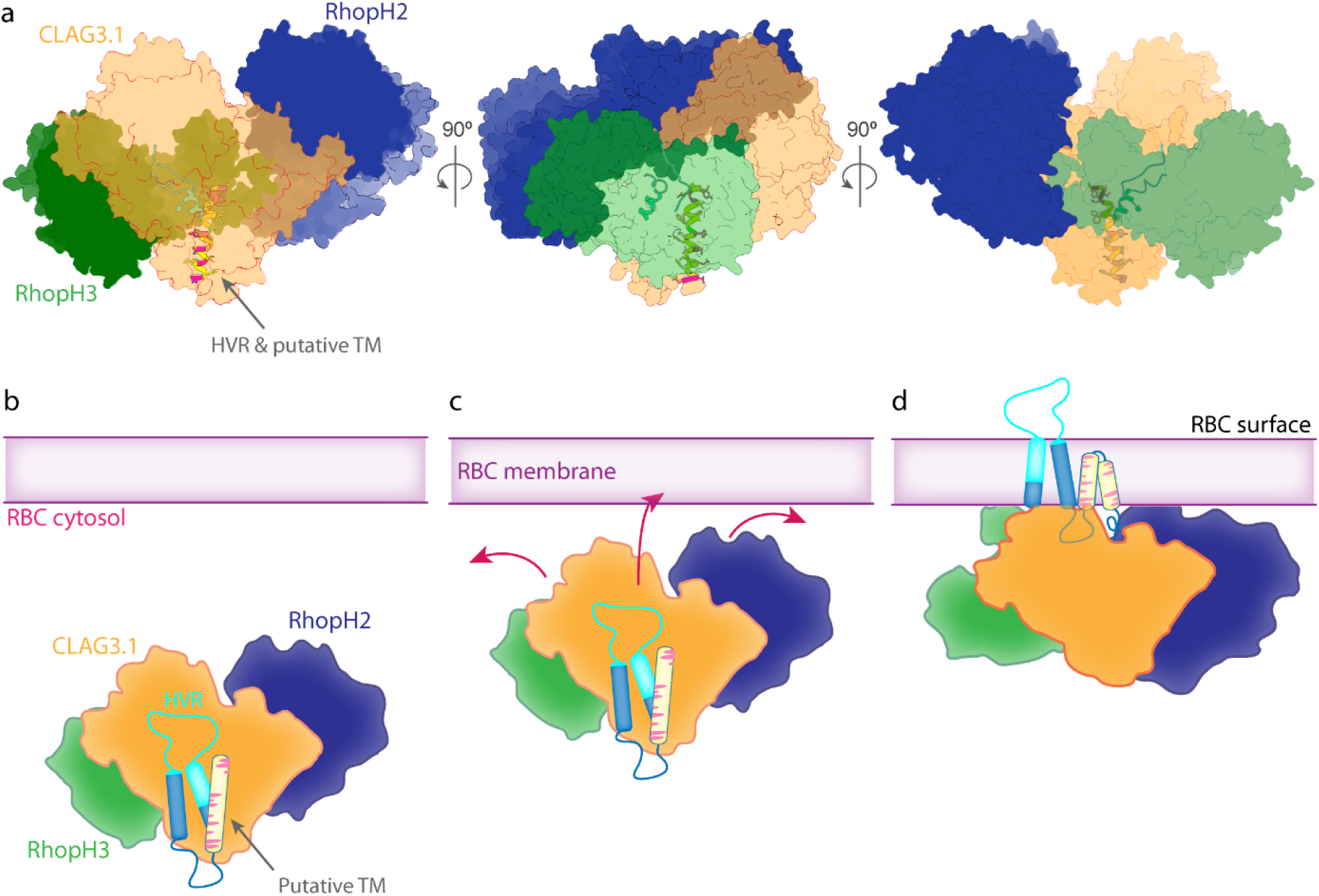
Model of CLAG3.1 putative transmembrane domain and HVR insertion into the erythrocyte membrane. **a,** Multiple views of the RhopH complex showing how the putative transmembrane and extracellular elements are buried in the core of the complex. The RhopH complex is displayed in space-filling surface diagram, with the putative transmembrane helix and HVR shown as ribbon diagrams. **b,** Schematic illustrating a model for how a conformational change in the RhopH complex may drive insertion of the extracellular and putative transmembrane elements into the red blood cell plasma membrane.

## Discussion

High-resolution structures provide invaluable frameworks for interpreting the wealth of biochemical and genetic data surrounding protein complexes implicated in the pathogenesis of malaria parasites^29^. Unfortunately, structural studies in malaria have been notoriously difficult^30^. Many *P. falciparum* proteins are extremely challenging to over-express recombinantly, with multi-protein complexes presenting a particular challenge due to the difficulty of recapitulating proper folding and assembly via reconstitution or co-expression in heterologous systems. Using our novel endogenous structural proteomics approach^21^, we circumvented these obstacles to determine the first near-atomic structure of the RhopH complex, enriched directly from parasite-infected red blood cells. Our discovery of a hypothesized—but hitherto unobserved—soluble state of the native RhopH complex is the first direct observation of an exported *P. falciparum* transmembrane protein in its soluble, trafficking state. This represents a major step forward in addressing the longstanding question of how parasite effectors, many of which are integral membrane proteins^4,5^, are trafficked to their final site of action in the erythrocyte membrane.

The structure provides exciting new insights into how trafficking works, revealing that previously predicted transmembrane and extracellular elements are buried in the middle of the complex, likely to protect these elements from the aqueous cytosolic environment during transit. Taken into account with all the data we have thus far, this discovery seems to point toward a model in which CLAG3, RhopH2, and RhopH3 associate early in the secretory pathway^8–10,12^, coming apart only for translocation across the parasitophorous vacuole membrane through PTEX^9,31^, and then reforming the soluble complex state in the erythrocyte cytosol to complete the journey out to the erythrocyte surface^9^. This suggests that the RhopH complex shares the ability of pathogenic pore-forming proteins to transition from a soluble form for traversing aqueous cytosolic environments to an integral transmembrane protein form upon arriving at its target membrane^27,32,33^, and that such transition may represent a general strategy employed by the parasite to deliver protein complexes to the erythrocyte membrane.

The ability to transition from a soluble form for traversing aqueous cytosolic or extracellular environments to an integral transmembrane protein form upon arriving at the target membrane is the unifying key feature shared by pore-forming proteins in biological systems ranging from bacteria to vertebrates^33^. As such, there is an argument for considering the RhopH complex to be a pore-forming protein complex. While the use of pore-forming proteins to unrestrictively permeabilize and lyse host-cell membranes during invasion and egress is a well-studied strategy commonly used by a wide variety of pathogens, including *P. falciparum* itself^27,33–35^, the use of pore-forming proteins to selectively alter host cell permeability to particular ions or solutes without destroying the membrane is less common and less well-understood. This strategy is rare in bacteria but essential for the function of several enveloped viruses, including influenza and human immunodeficiency virus (HIV)^33^. In this respect, *P. falciparum*’s use of the RhopH/PSAC complex to modulate host erythrocyte membrane permeability for nutrient acquisition is more similar to how viruses use viroporins to establish ion-specific permeability across host-cell membranes, making the RhopH complex distinct from the more common use of pore-forming proteins by malaria parasites to perforate host cell membranes during invasion and egress^35^. Similar to most viroporins^32^, the vast majority of the RhopH complex is located intracellularly, with only a small segment exposed on the erythrocyte membrane surface. The mechanisms underlying viroporins’ transitions from soluble to membrane-bound, often triggered by a change in pH or interaction with a lipid or protein in the target membrane, may hold hints about the possible mechanism by which elements of CLAG3.1 are inserted into the erythrocyte membrane. Given the unusual abundance of both cysteines and disulfide bonds in the RhopH complex, we speculated that the large conformational change the RhopH complex would need to undergo at the membrane to expose and insert the putative transmembrane and extracellular elements into the erythrocyte membrane might be triggered or regulated by allosteric disulfide bonds.

First discovered more than a decade ago, the formation and breaking of allosteric disulfide bonds have been firmly established as a strategy used in wide range of biological systems for regulating and modulating protein structure and function^36,37^. Indeed, when we examined the 14 disulfide bonds in our structure, we found three that exhibited the well-defined strained geometry that has been found to be the hallmark of all allosteric disulfide bonds characterized thus far (Supplementary Table 5)^36^ – one in RhopH2, and two in RhopH3 (Supplementary Fig. 3). As such, it is possible that these bonds may break or be cleaved upon interaction with the erythrocyte membrane, triggering a conformation change that extrudes the putative transmembrane and extracellular elements from the middle of the complex and drives them through membrane. While no parasite-derived protein disulfide isomerases (PDIs) have been identified in the *P. falciparum* exportome, human erythrocytes do carry active PDIs in the plasma membrane^38,39^, which could cleave the allosteric disulfides and trigger the conformational change upon coming in contact with the RhopH complex at the membrane. Alternatively, different redox potential in the cytosol could trigger the breakage of the disulfide bonds to effect the massive conformational changes needed to expose and bring the above-described helical bundle of CLAG3 to interact with red blood cell membrane (Fig. 5b).

Although surface-exposed membrane proteins with essential functions in pathogenesis are often popular targets for drug development, these proteins have thus far proven to be difficult targets for drug and vaccine development in *P. falciparum*, due to various mechanisms the parasite employs to evade immune detection. If the use of a soluble form is a general strategy employed by the parasite to deliver proteins to the erythrocyte membrane, high resolution structures of the soluble trafficking forms of these essential membrane protein complexes present an opportunity to approach development of therapies from a new angle, focused on blocking assembly or trafficking of complexes before they have a chance to insert into the erythrocyte membrane. As such, the novel approach presented here for overcoming the barriers to high-resolution structural studies in malaria parasites opens the door for developing new anti-malarial therapies with novel modes of action and can also be applied to hard-to-isolate complexes in other biological systems to answer longstanding questions.

## Acknowledgements

This research was supported in part by grants from National Institutes of Health (R01GM071940/AI094386/DE025567 to Z.H.Z. and K99/R00 HL133453 to J.R.B.). CM.H. acknowledges funding from the Ruth L. Kirschstein National Research Service Award (AI007323). We thank the UCLA Proteome Research Center for assistance in mass spectrometry and acknowledge the use of resources in the Electron Imaging Center for Nanomachines supported by UCLA and grants from NIH (S10RR23057, S10OD018111 and U24GM116792) and NSF (DBI-1338135 and DMR-1548924).

## Author Contributions

CMH initiated the project; JRB cultured and harvested parasite material; CMH purified the sample from parasite pellets, screened purified samples by negative stain, optimized sample freezing conditions for cryoEM, acquired and processed the cryoEM data, identified the components of the RhopH complex in the cryoEM density map using cryoID, interpreted the structures, and wrote the paper; ML and JJ built and refined the atomic models and JJ helped interpret the structures; XL helped with the initial identification of CLAG3.1 from the cryoEM density map using cryoID. ZHZ supervised the cryoEM aspects of the project, interpreted the structures and wrote the paper; DEG supervised parasitology aspects of the project. JJ, JRB, and DEG helped edit the paper; all authors approved the paper.

## Data Availability

The atomic models and the cryoEM density maps are deposited to the Protein Data Bank and the Electron Microscopy Data Bank, under the accession numbers of XXXX, EMD-XXXXX, respectively.

## Materials and Methods

### Parasite culture

*P. falciparum* cultures were prepared as described previously^21,29^. Erythrocytes were then lysed with 0.0125% saponin (Sigma, sapogenin content ≥ 10%) in cold phosphate-buffered saline (PBS) with an EDTA-free protease inhibitor cocktail (Roche or Pierce). The released P. falciparum parasites were then washed in cold PBS with EDTA-free protease inhibitor cocktail. The washed cell pellets were flash-frozen in liquid nitrogen and stored at −80 °C.

### Sucrose gradient fractionation *P. falciparum* parasite lysate and subsequent evaluation of fractions using negative stain electron microscopy

Frozen parasite pellets were resuspended in lysis buffer (25mM HEPES pH7.4, 150mM KCl, 10mM MgCl_2_, 10% glycerol) and lysed using a glass Dounce tissue homogenizer. The soluble lysates were separated from the membrane fraction by centrifuging at 100,000g for 1 h and then fractionated on a sucrose gradient as previously described^21^. The resulting fractions were evaluated for the presence and relative abundance of proteins of interest as described previously^21^. Briefly, fractions were assessed for the abundance of potential particles of interest based on silver-stained SDS-PAGE, tryptic digest liquid chromatography mass spectrometry, and negative stain electron microscopy. The conventional approach of using gel filtration chromatography to assess sample quality was eschewed in favor of using negative stain electron microscopy to directly visualize and evaluate the abundance of intact particles giving rise to promising 2D class averages with distinct features. As such, small datasets of ~100,000 particles were collected for each fraction and 2D class averages generated in RELION^22,40^ were used to identify fractions containing promising particles (Fig. 1e).

### Mass spectrometry

Mass spectrometry was performed as previously described^21^. The proteomics data will be deposited to the ProteomeXchange Consortium (http://proteomecentral.proteomexchange.org) via the MassIVE partner repository with the dataset identifier XXXXXXXXX.

### Cryo electron microscopy

CryoEM grids of the selected fraction of parasite lysate were prepared and imaged as described previously^21^, yielding a final dataset consisting of a total of 19,449 movies.

### Image processing and 3D reconstruction

Frames in each movie were aligned, gain reference-corrected and dose-weighted to generate a micrograph using MotionCor2 [^41^]. Micrographs that were aligned without dose-weighting were also generated for contrast transfer function (CTF) estimation using CTFFIND4^42^ and automated particle picking in Gautomatch^43^.

2,900,167 particles were extracted from 19,449 micrographs. Two to three rounds of reference-free two-dimensional (2D) classification in RELION were used to exclude junk particles. 858,049 particles belonging to 2D class averages exhibiting clear secondary structure features were further classified in CryoSPARC^23^. A subset of the 2D class averages, corresponding to 108,872 particles, were identified as likely to have originated from the same asymmetric volume. These 108,872 particles were subjected to an unsupervised single-class *ab initio* 3D reconstruction followed by a single-class homogenous refinement using C1 symmetry in CryoSPARC. The resulting reconstruction was then subjected to non-uniform refinement using C1 symmetry in CryoSPARC, yielding a density map with a final overall resolution of 3.72Å (Fig. 2c-d, Supplementary Fig. 5). The 108,872 particles were then subjected to focused classification and refinement in RELION to reach sufficient local resolutions throughout the map to allow successful identification of all three proteins in the map using cryoID.

### Identifying proteins in the cryoEM density map using cryoID

We ran the cryoID *Get_queries* subprogram ^21^ on the initial 3.7Å resolution density map obtained from refinement and postprocessing in RELION, using default symmetry (Any) and a high resolution limit of 3.6Å as the input parameters. We then manually inspected the resulting query models, correcting residues incorrectly assigned by *Get_queries* and extending the queries on both ends as the density permitted (Fig. 2e). This yielded the following degenerate sequences, which were then used for searching:

***Query Set 1:***

1. GLGYGYYGGLGYYGGGLYYYLLXYYG
2. LLGGPPGGXGYLYLLGLGLGYYYXLYLL-GXGGYYYXXY

Using this set of query sequences, *cryoID* identified two candidates for the protein in this region of the density map from a candidate pool consisting of the 1170 proteins identified in this sucrose gradient fraction by mass spectrometry. The two candidates, CLAG3.1 and CLAG3.2 from *P. falciparum* (O77309/ O77310), share 96% protein sequence identity (Supplementary Table 2). We confirmed the identification by manually building a *de novo* atomic model into the rest of the map (Fig. 2b). Focused classification and refinement was used to improve the local resolution around the region corresponding to residues 1110-1150, one of the few segments in which CLAG3.1 and CLAG3.2 differ in sequence. The improved resolution was sufficient for us to confidently determine that the sequence in this region matches CLAG3.1 rather than CLAG3.2 based on the presence of two residues, T1140 and H1141, which are found in CLAG3.1 but not CLAG3.2, confirming that the protein in the complex is CLAG3.1 (Supplementary Fig. 1).

We completed a backbone trace of the remaining regions of the density map that did not correspond to CLAG3.1, which allowed us to determine that there were likely two additional, distinct polypeptide chains represented in the density map. We then used *Get_queries* to generate queries from these two remaining regions of the density map, yielding two additional sets of queries (Fig. 2f-g). We manually inspected the query models, correcting residues incorrectly assigned by *Get_queries* and extending the queries on both ends as the density permitted. This yielded the following degenerate sequences, which were then used for searching:

***Query Set 2:***

1. GLYLKKGGGGWGYLLKGLGLYGYLLGLLG
2. GLGLGLGYGYGLGGKL
3. GYGLGYLLYLGLGGLL

***Query Set 3:***

1. GYYGGGLLLGGYGLGYYLGKYLL
2. YGKGGGGGGLYGGGGLY

Using these two sets of query sequences, *cryoID* identified the proteins in the two regions of the density map, from a candidate pool consisting of the 1170 proteins identified in this sucrose gradient fraction by mass spectrometry, to be RhopH2 (Query set 2) and RhopH3 (Query set 3) from *P. falciparum* (C0H571 and Q8I395) (Supplementary Table 3,4). We subsequently confirmed the identification by manually building *de novo* atomic models into the corresponding two regions of the map (Fig. 2b).

### Manual model building and refinement

Map interpretation was performed with UCSF Chimera^44^ and COOT^45^. *P. falciparum* protein sequences were obtained from the National Center for Biotechnology Information (NCBI)^46^ and the PlasmoDB^47^ protein databases. Initial sequence registrations during model building for all three proteins in the map were determined using cryoID^21^. PHYRE2^48^ secondary structure predictions were also used as a guide during subsequent model building. Each residue in the three proteins was manually traced and built *de novo* in COOT, using the best focused classification map for each local area. Rotamers for residues were manually selected as assessed by residue-to-map fit using the select_rotamer option in COOT.

Map resolution outside of core regions in the complex were not sufficient to permit the use of real_space_refine_zone found in COOT. As such, manual refinement targeting both protein geometry and fit with the density map was carried out using COOT’s regularize_zone and fixed_atoms_for_refinement functions. Finally, resulting models for the complexes were subjected to iterative cycles of automated refinement using the phenix.real_space_refine program in PHENIX^49^ followed by further manual refinement, performed against a map visually determined to possess the best mix of overall features for each local region, to achieve the final structure.

All figures and videos were prepared with UCSF Chimera, Pymol^50^, and Resmap^51^. Molprobity^52^ was used to validate the stereochemistry of the final models.

## Supplementary Figures

**Supplementary Figure 1.**
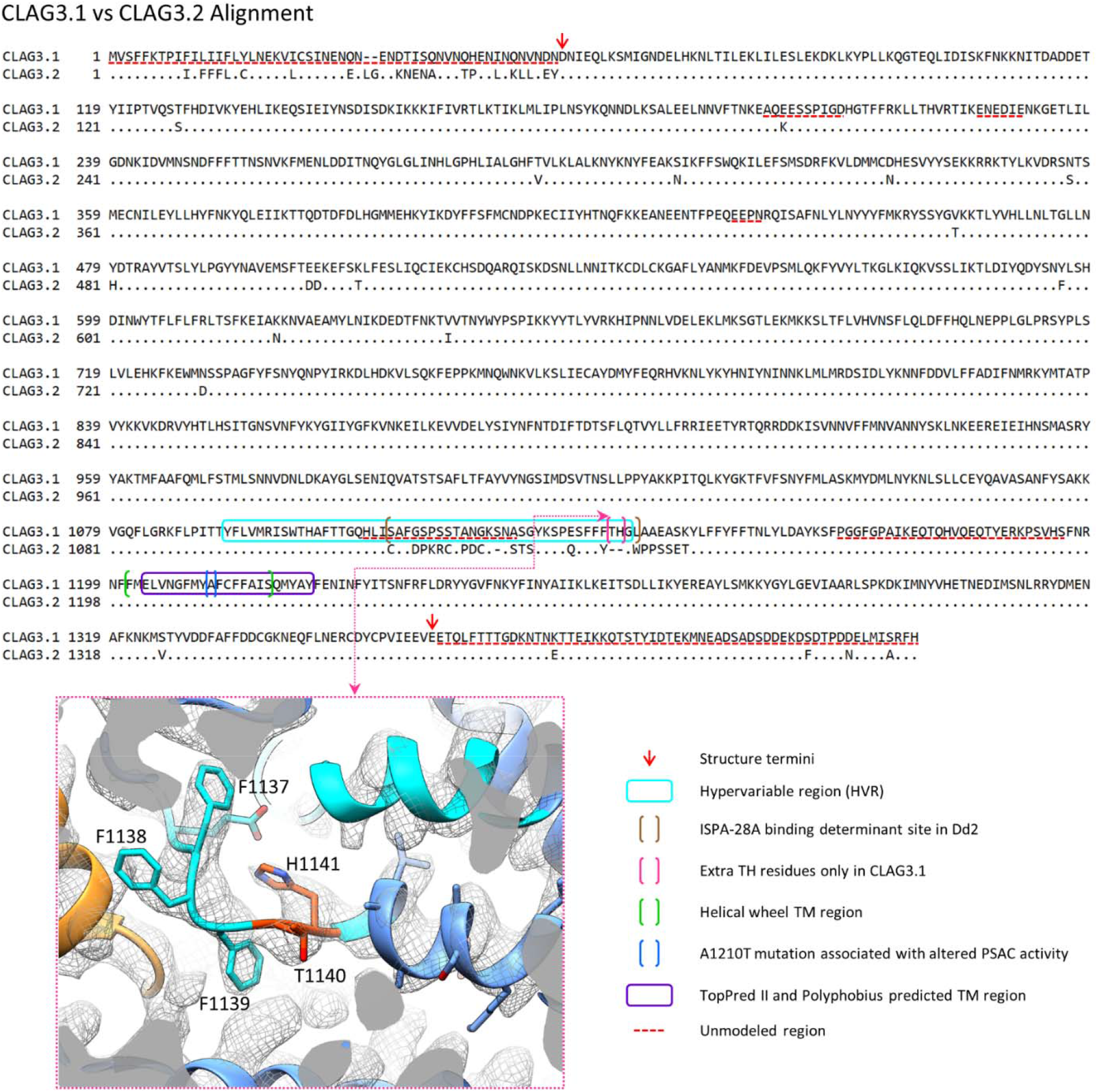
CLAG3.1 vs CLAG3.2 alignment. Annotated alignment of CLAG3.1 vs CLAG3.2, with periods (.) denoting consensus residues in CLAG3.2. Key denotes notable CLAG3.1 features. Inset shows the a.a. 1136-1148 region of CLAG3.1 depicted in ribbon/ball-and-stick model and density mesh. T1140 and H1141 (orange residues) were critical in identifying the CLAG variant within our RhopH complex as CLAG3.1.

**Supplementary Figure 2.**
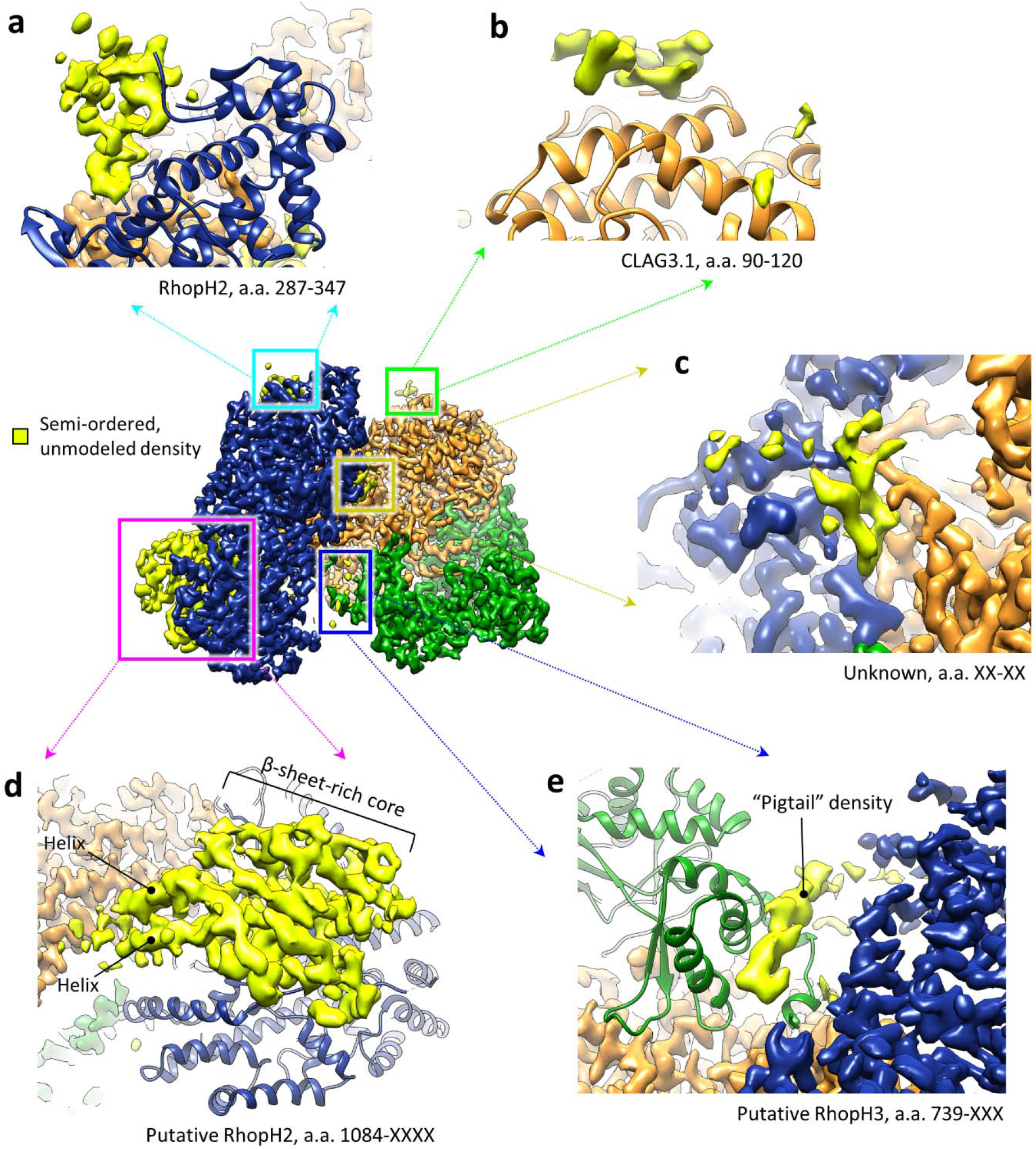
Unmodeled semi-ordered density in the RhopH complex. **a-e,** Regions of unmodeled semi-ordered density observable in the RhopH complex are depicted in neon yellow, along with descriptions of assessments of their identity.

**Supplementary Figure 3.**
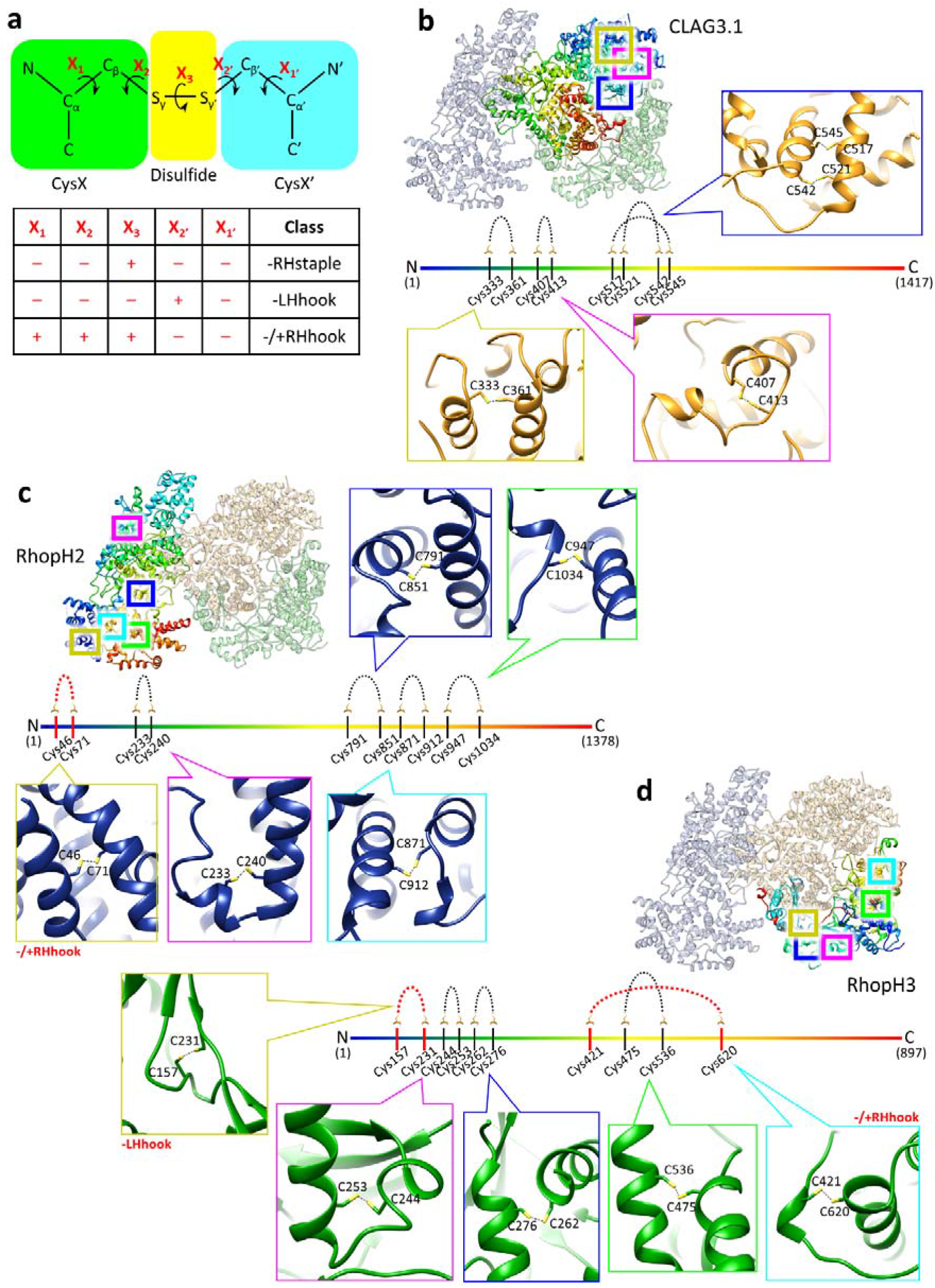
Disulfide bridges of the RhopH complex. **a,** Graphic depicting the Χ angles (red) of a disulfide bond. Chart displays the signs, positive (+) or negative (–), of sets of Χ angles in a disulfide bond that define the three classes often associated with allosteric disulfides. **b-d,** Disulfide bridges observed in CLAG3.1 (**b**), RhopH2 (**c**), and RhopH3 (**d**) mapped onto each protein’s primary sequence. Respective insets show close-up ribbon/ball- and-stick representations of cysteine-cysteine interactions. Disulfides displaying Χ angles of the three allosteric disulfide classes are marked with red graphics.

**Supplementary Figure 4.**
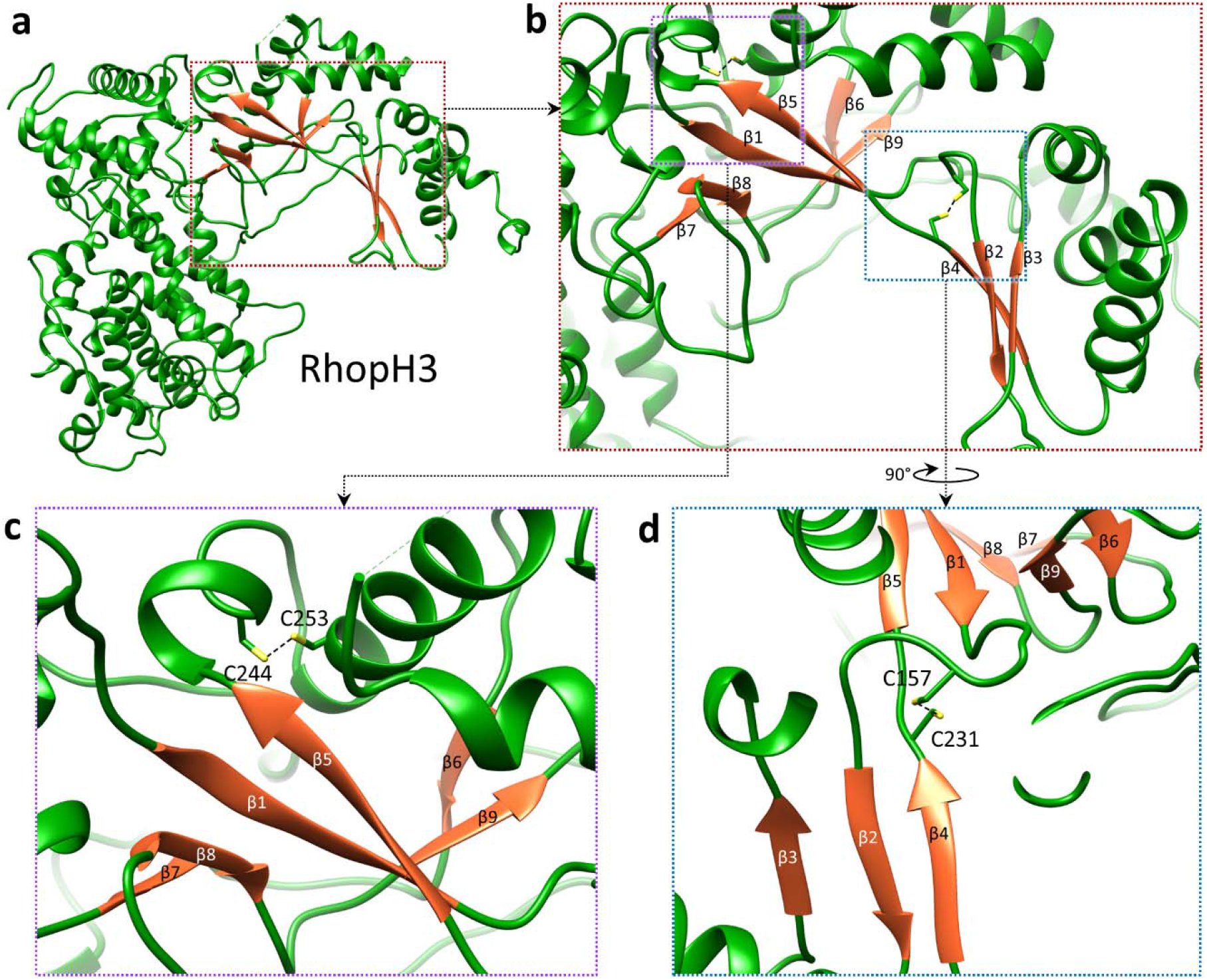
Cysteine interactions in RhopH3’s β-sheet motifs. **a-b,** RhopH3 contains three sets of β-sheets (β-strands colored orange) that comprise an interleaved β-sheet motif. **c-d,** Two prominent disulfide bridges between C244 and C253 (**c**) and C157 and C231 (**d**) appear to stabilize a continuous peptide sequence (a.a. 225-245), containing β4 and β5, that is a core part of the motif.

**Supplementary Figure 5.**
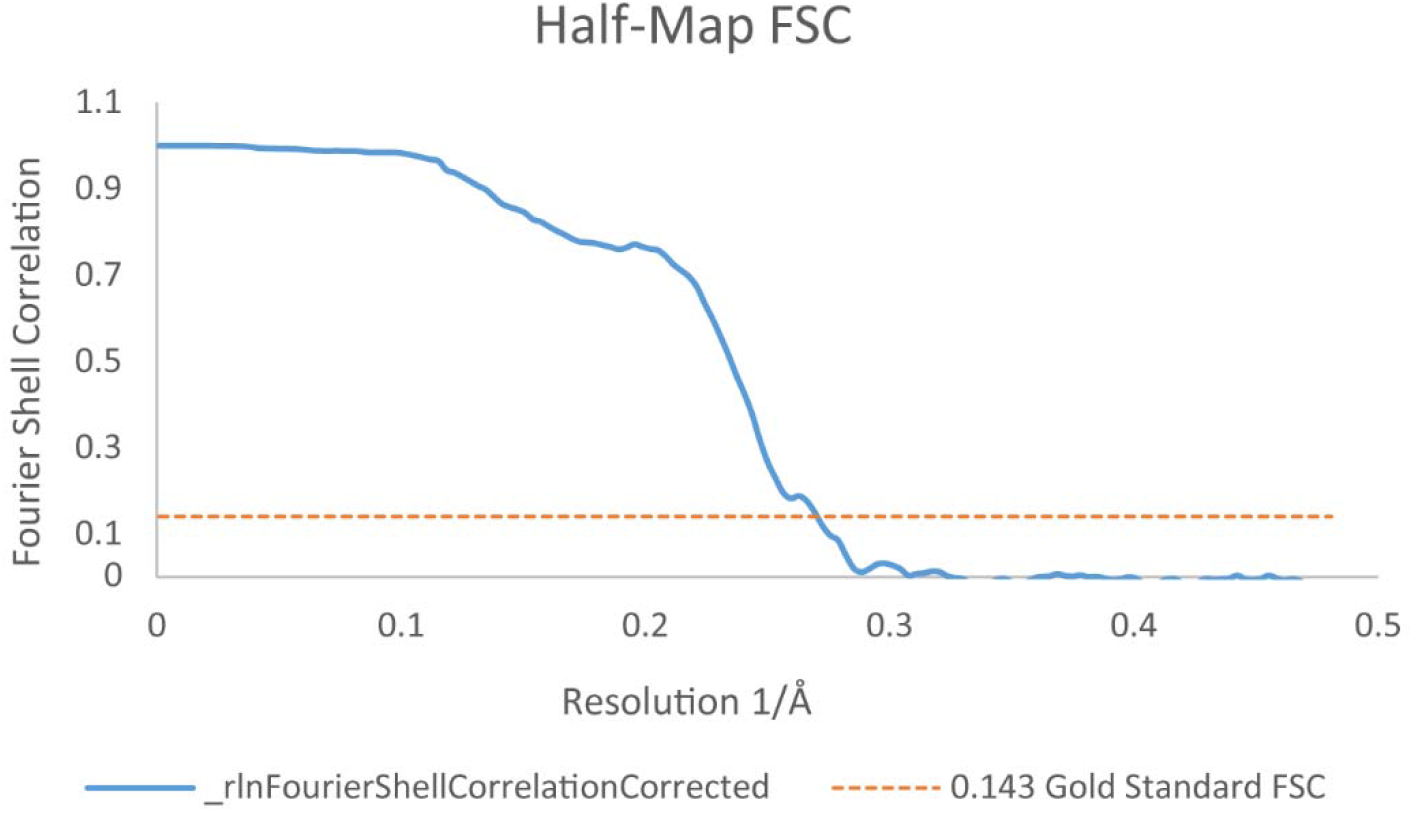
Half-map FSC of the RhopH complex structure.

**Supplementary Table 1 | Mass Spectrometry Data – Uploaded as an Excel File**

**Supplementary Table 2.**
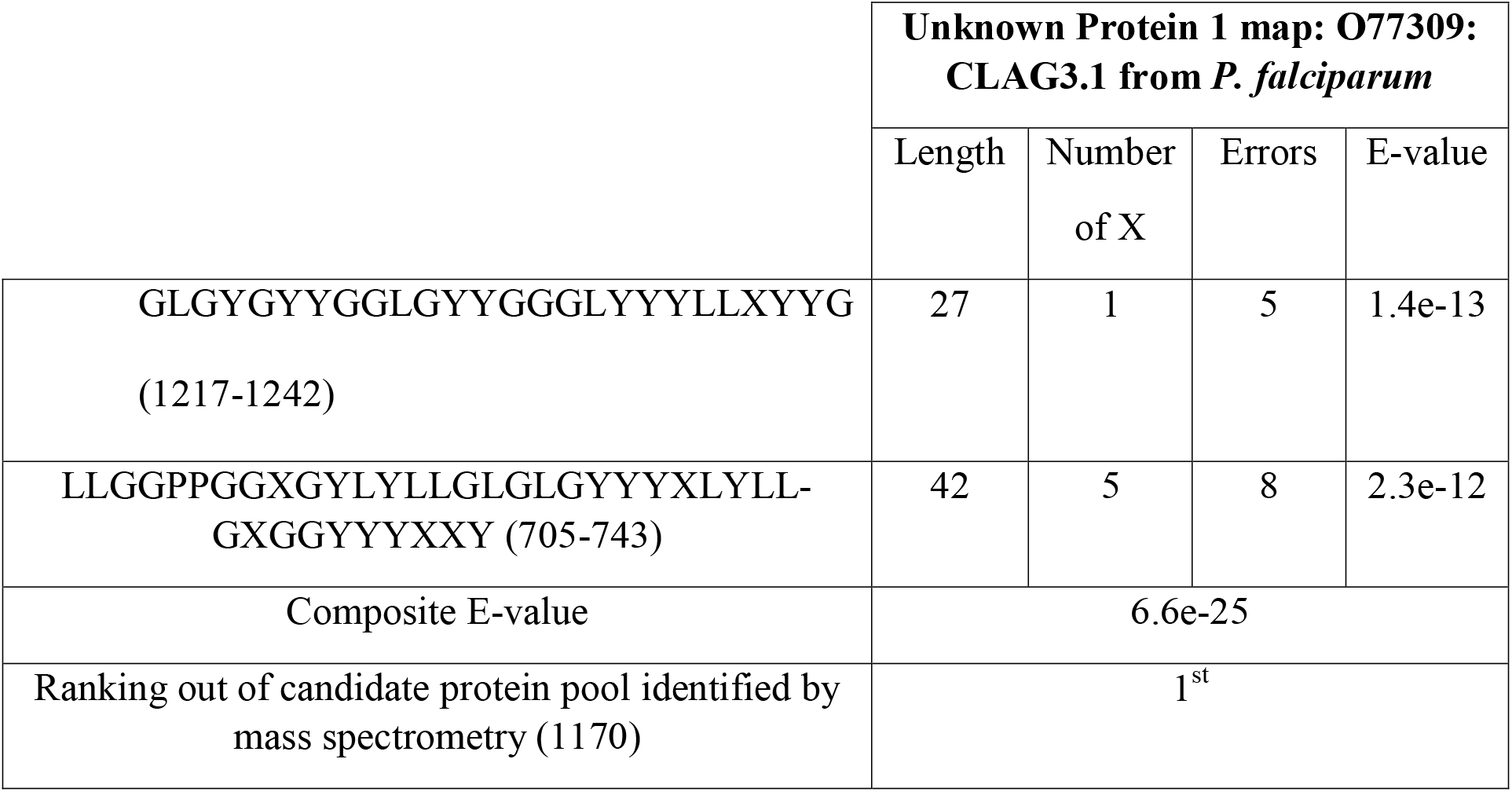
Unknown Protein Complex Component 1 *cryoID* results.

**Supplementary Table 3.**
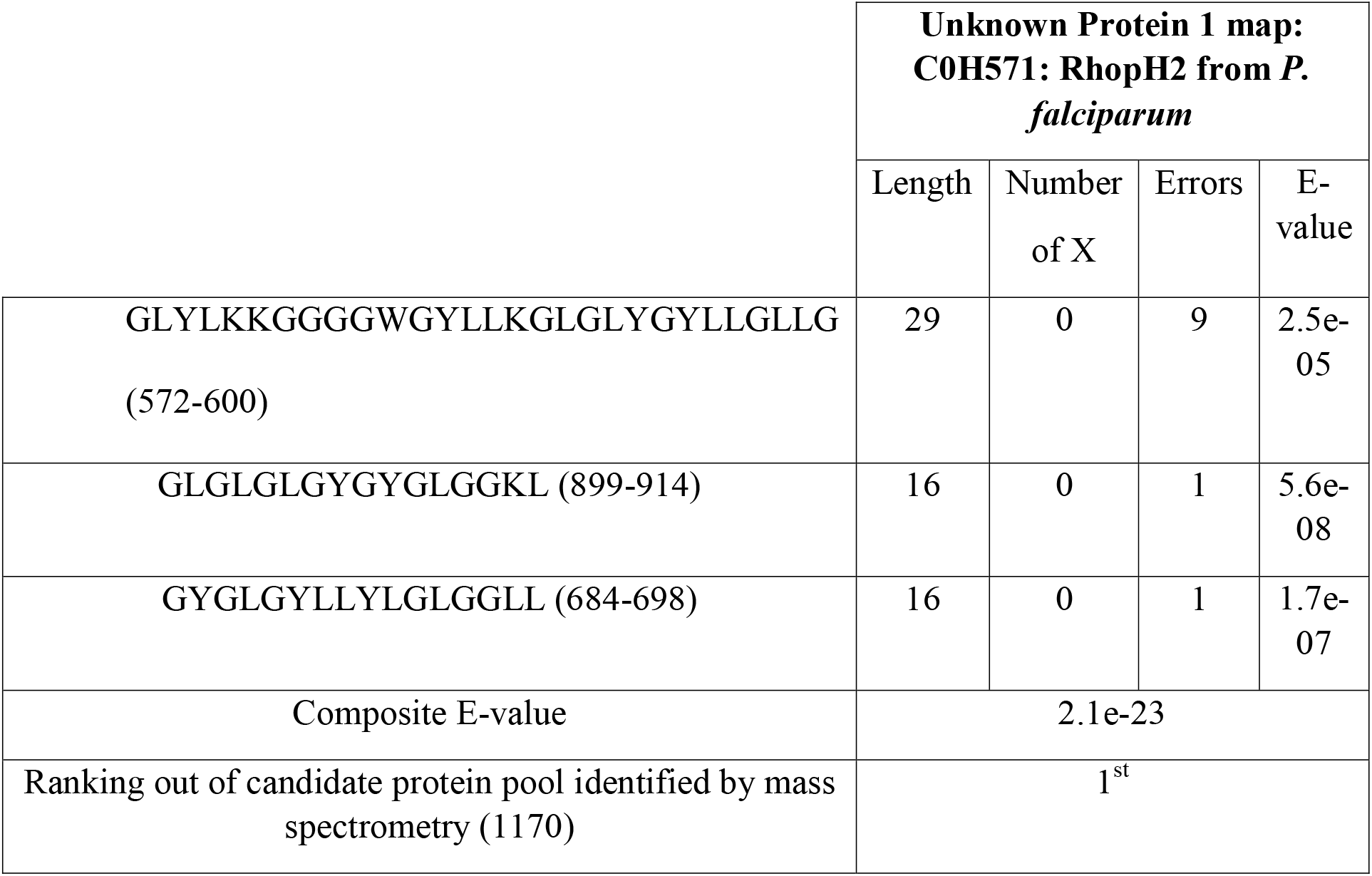
Unknown Protein Complex Component 2 *cryoID* results.

**Supplementary Table 4.**
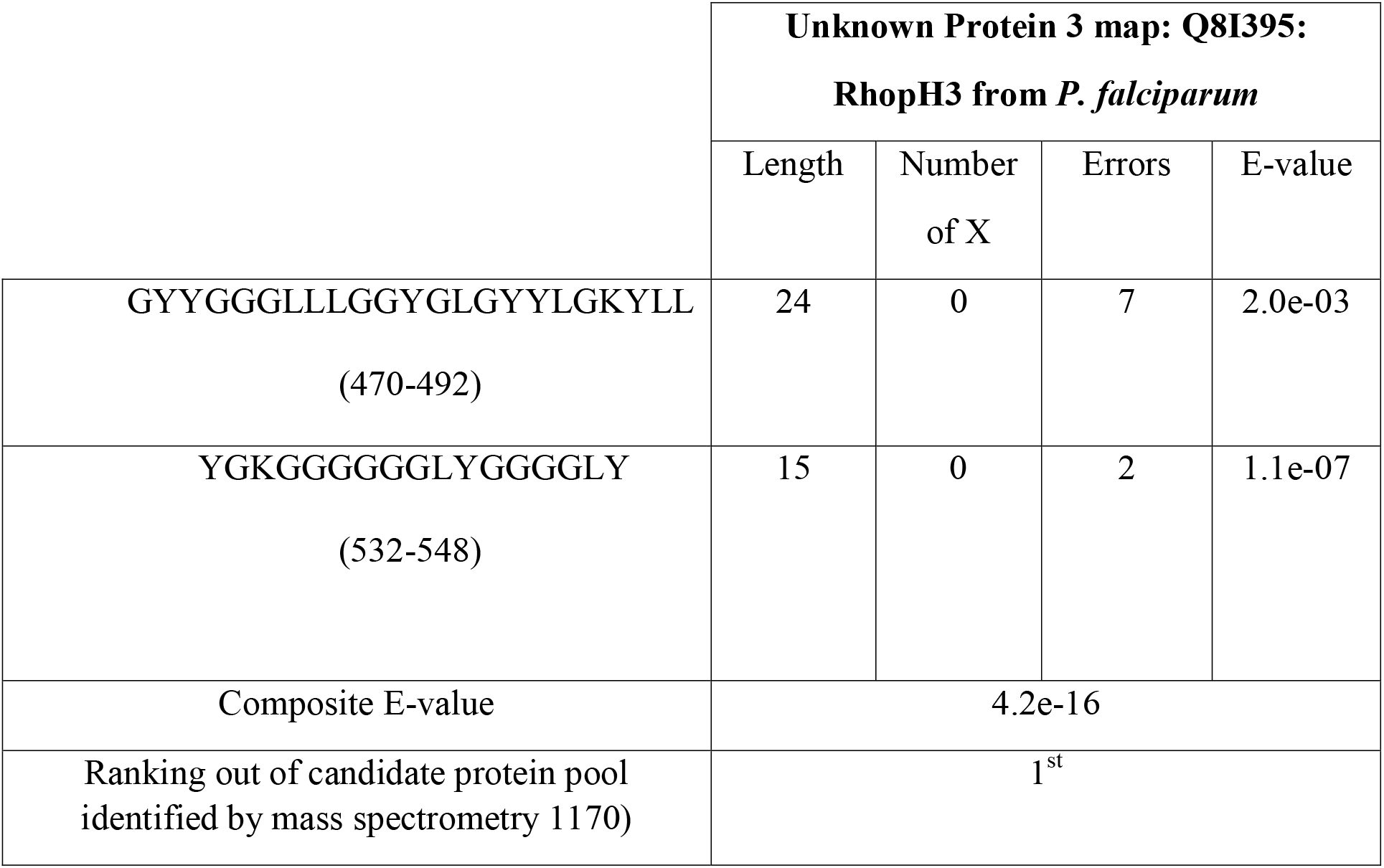
Unknown Protein Complex Component 3 *cryoID* results.

**Supplementary Table 5.** RhopH complex disulfide Χ angle analysis. Red values denote disulfide classifications associated with allosteric disulfide bonds.

|  |  |  |  |  |  | DSE (kJ/mol) | Classification |
| --- | --- | --- | --- | --- | --- | --- | --- |
| X angles | X<sub>1</sub> | X<sub>2</sub> | X<sub>3</sub> | X<sub>2'</sub> | X<sub>1'</sub> |  |  |
| **CLAG3.1** |  |  |  |  |  |  |  |
| cys333-cys361 | -56.08 | 168.97 | 118.51 | 110.74 | -167.34 | 22.190535 | -RHspiral |
| cys407-cys413 | -65.38 | 128.45 | 92.12 | 174.46 | 67.56 | 11.938369 | +/-RHspiral |
| cys517-cys545 | 178.68 | -161.82 | 94.68 | 176.94 | -71.43 | 6.598143 | +/-RHhook |
| cys521-cys542 | 171.74 | 61.26 | 148.8 | -68 | -80.11 | 29.459694 | -/+RHhook |
| **RhopH2** |  |  |  |  |  |  |  |
| cys46-cys71 | -68.13 | -115.68 | 146.14 | 100.81 | 61.67 | 38.658783 | -/+RHhook |
| cys233-cys240 | 178.27 | 84.28 | -85.71 | -172.02 | -176 | 5.638452 | -/+LHhook |
| cys791-cys851 | 176.99 | -167.05 | -105.17 | -107.32 | 179.46 | 14.808307 | +LHspiral |
| cys871-cys912 | -175.71 | -142.04 | -72.85 | -90 | -177.01 | 13.459173 | -LHspiral |
| cys947-cys1034 | 178.3 | 50.74 | 106.83 | 1.75 | -166.31 | 17.817829 | +/-RHspiral |
| **RhopH3** |  |  |  |  |  |  |  |
| cys157-cys231 | -160.85 | 80.66 | -163.72 | -82.56 | -69.86 | 37.605534 | -LHhook |
| cys244-cys253 | -62.19 | -29.86 | -96.99 | -176.49 | -177.68 | 8.239258 | -LHspiral |
| cys262-cys276 | -162.47 | 33.06 | 66.46 | 66.49 | 178.95 | 11.862575 | +/-RHspiral |
| cys421-cys620 | -80.05 | -89.78 | 122.21 | 76.31 | 179.74 | 23.093697 | -/+RHhook |
| cys475-cys536 | 178.02 | 89.12 | 102.85 | 30.87 | -164.81 | 16.054579 | +/-RHspiral |

## Supplementary Movie Stills and Legends

**Supplementary Video 1.**
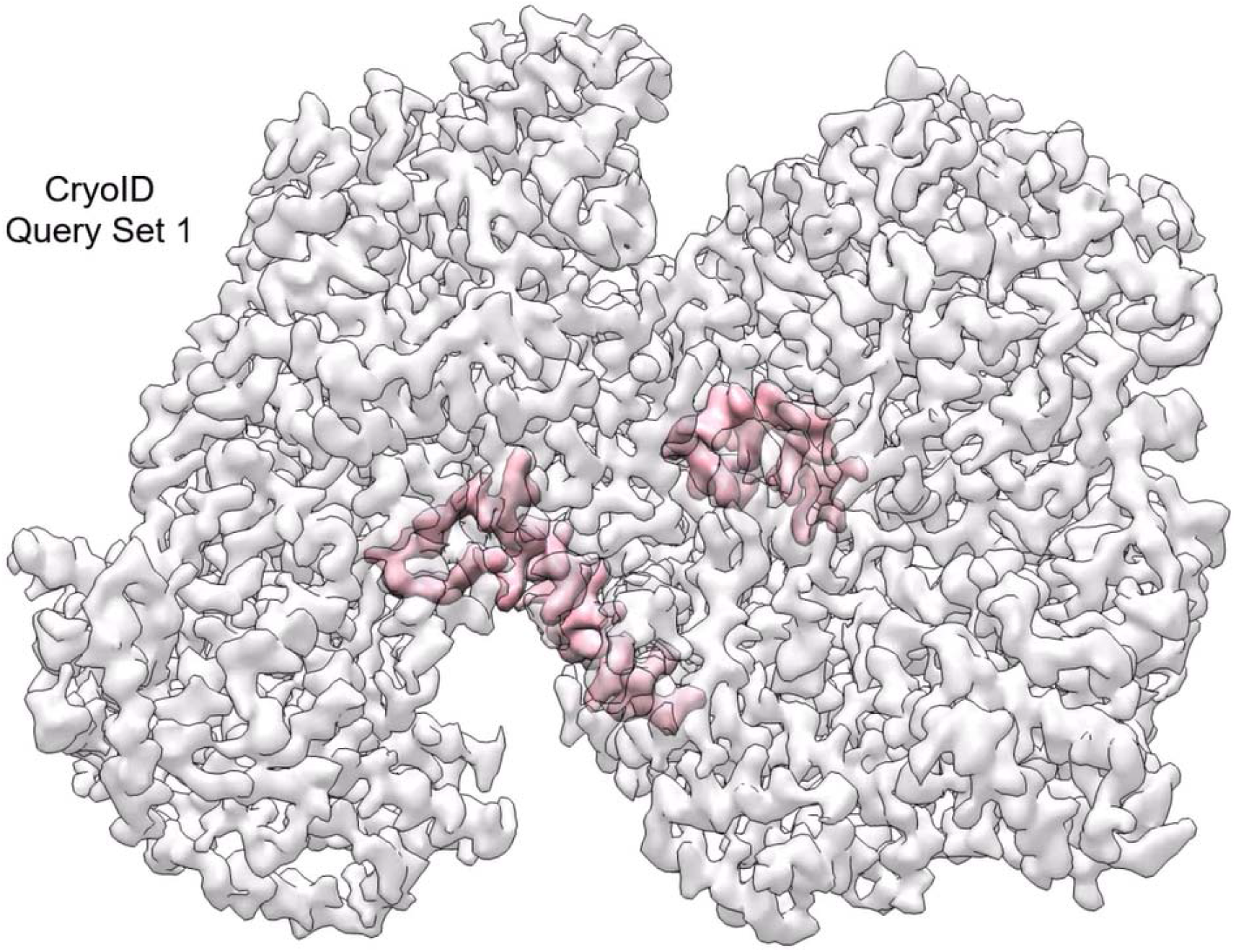
Movie highlighting the details of the cryoID query set used to identify CLAG3.1 from the cryoEM density map.

**Supplementary Video 2.**
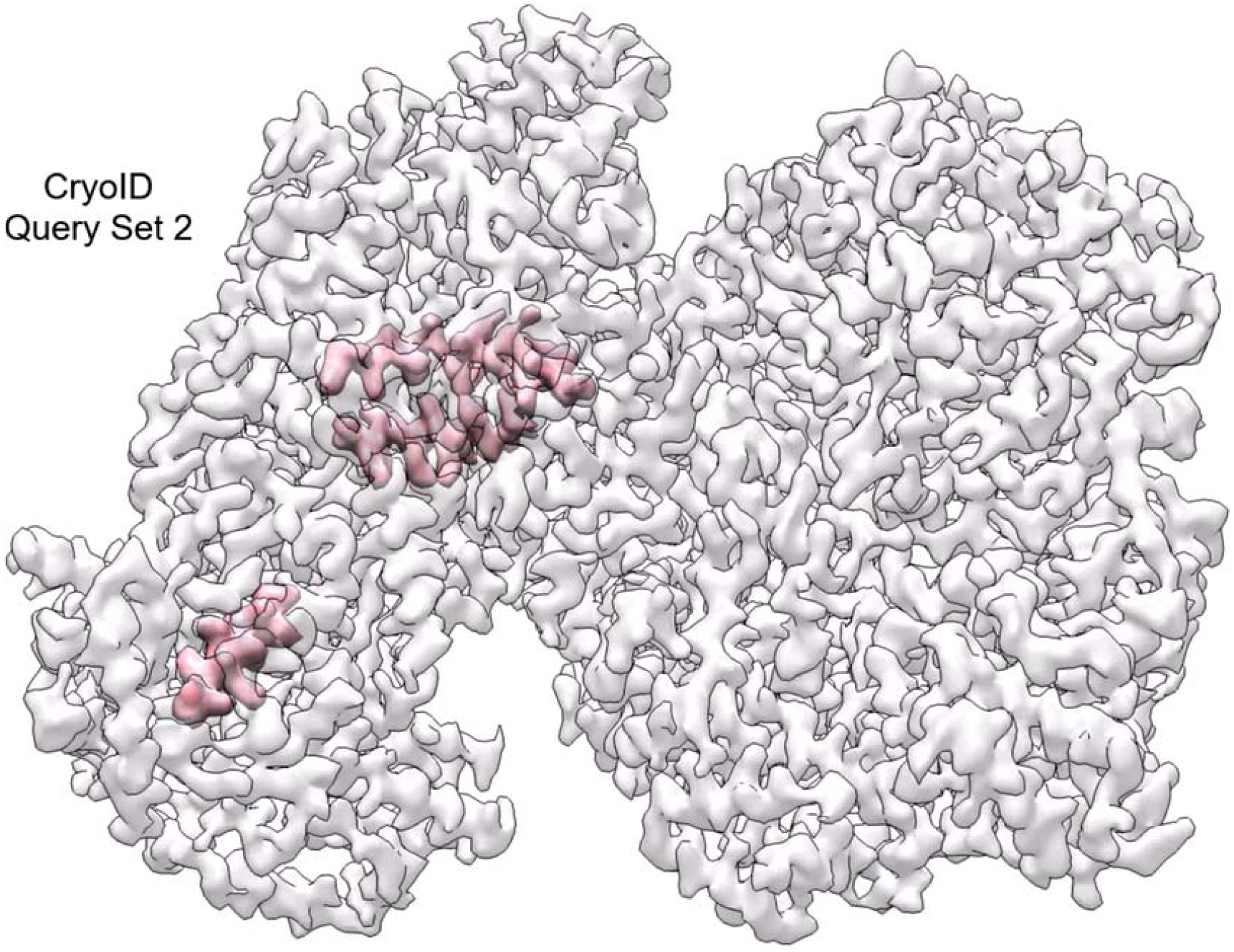
Movie highlighting the details of the cryoID query set used to identify RhopH2 from the cryoEM density map.

**Supplementary Video 3.**
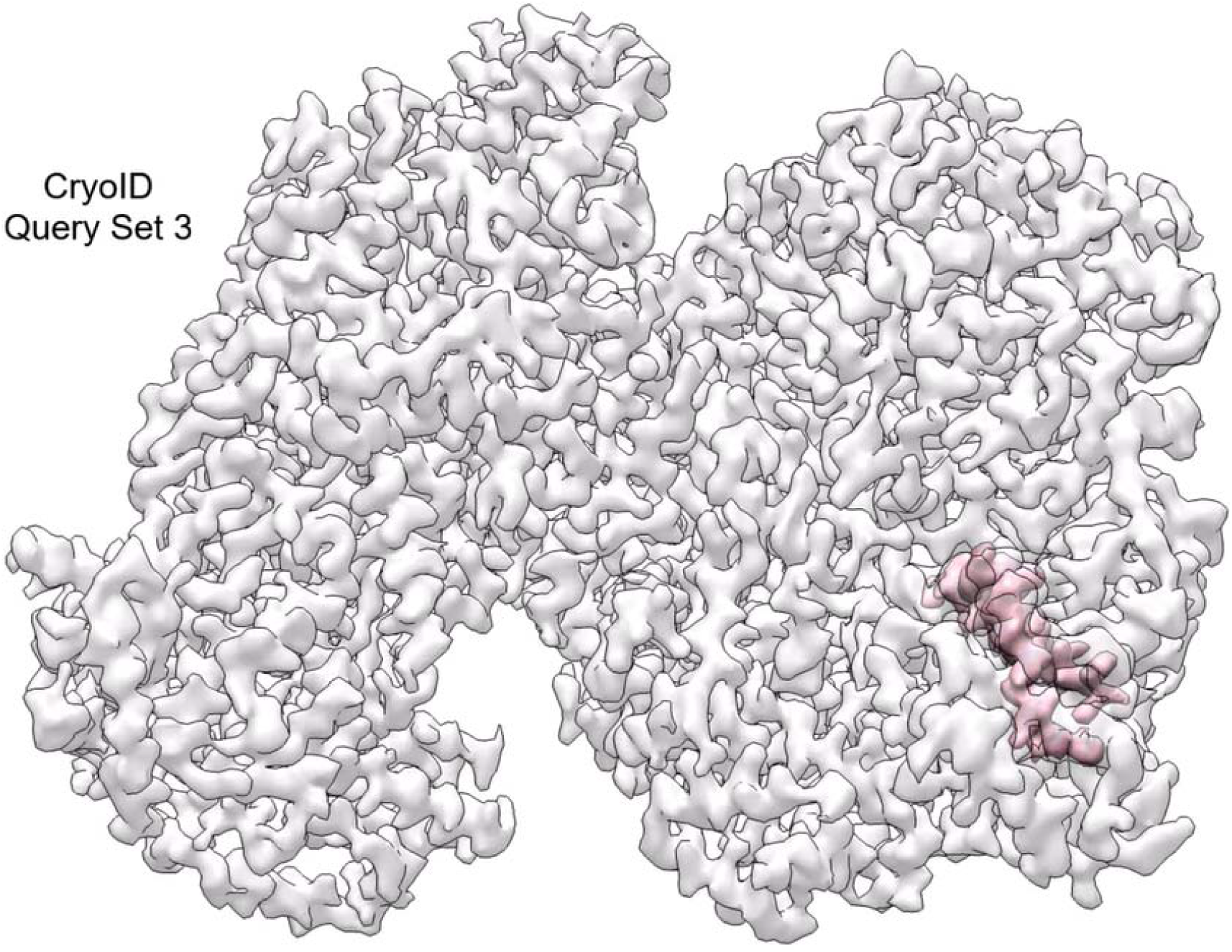
Movie highlighting the details of the cryoID query set used to identify RhopH3 from the cryoEM density map.

**Supplementary Video 4.**
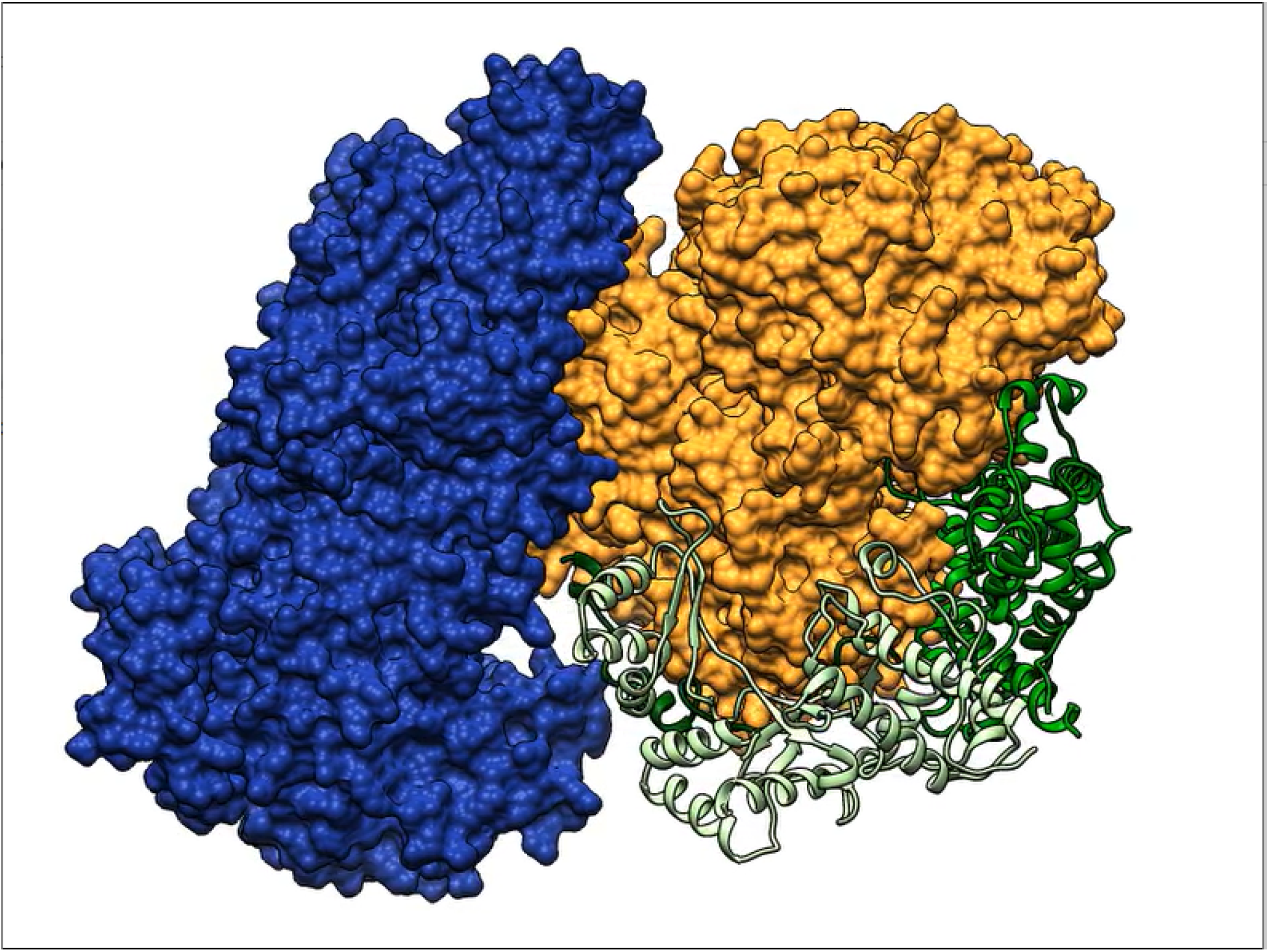
360° view of the RhopH complex displaying the interaction interfaces of each member of the RhopH complex with the other two components.

**Supplementary Video 5.**
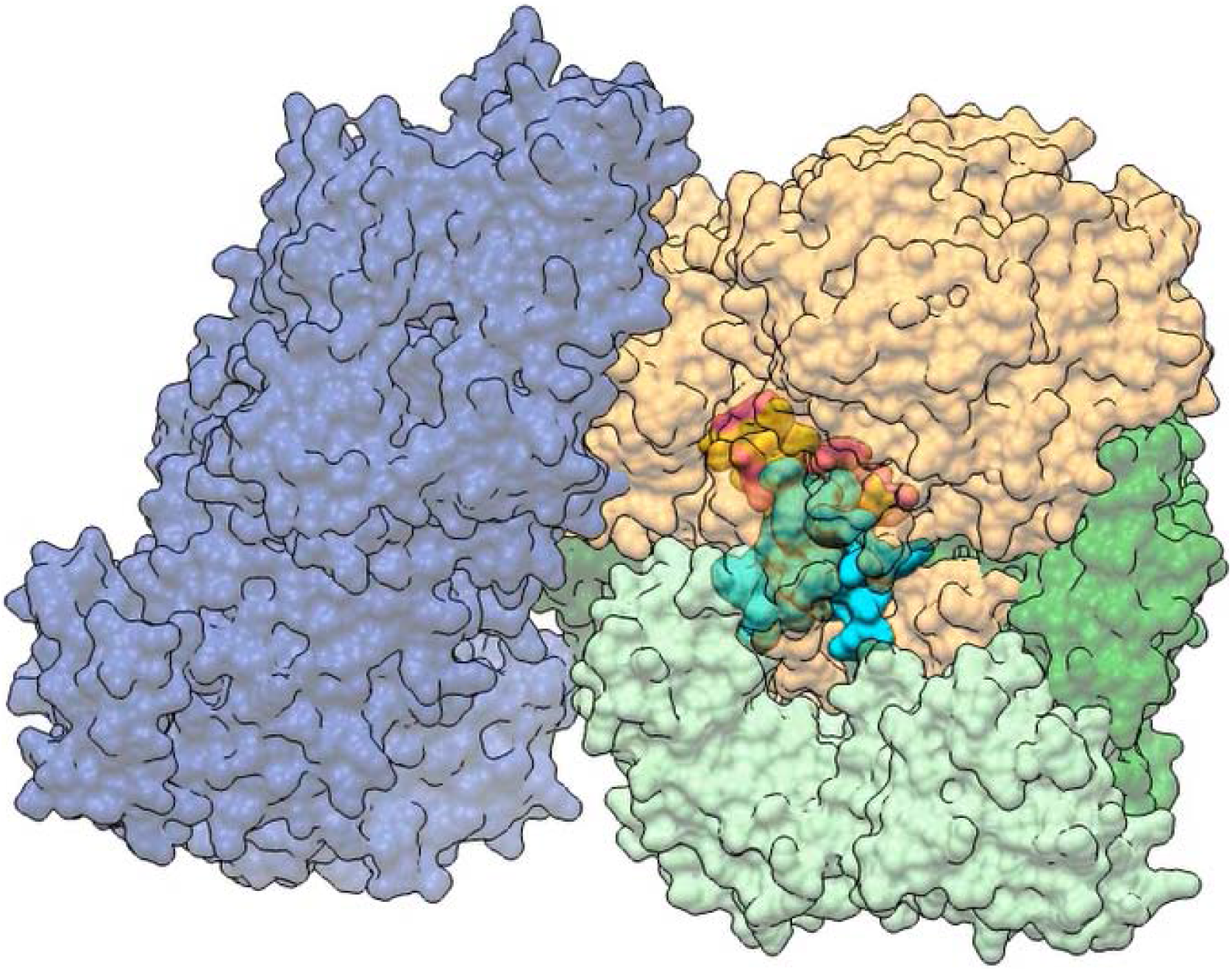
360° view of the RhopH complex, showing how the putative transmembrane helix (gold and pink) and extracellular hypervariable region (cyan) are embedded in the middle of the complex.

**Supplementary Video 6.**
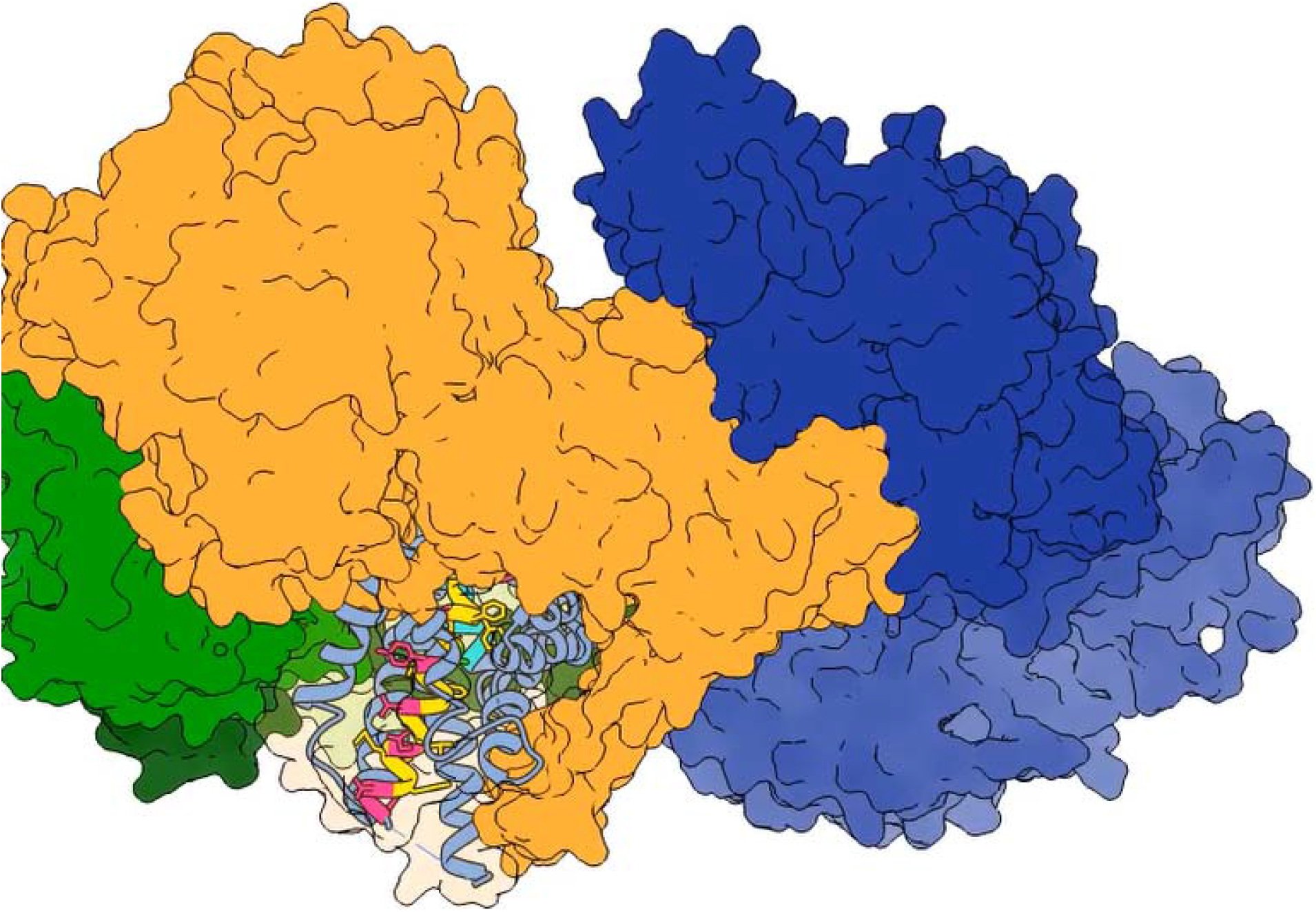
Detailed views of the CLAG3.1 helical bundle surrounding the putative transmembrane helix and extracellular hypervariable region of CLAG3.1, buried inside the RhopH complex.

## References

1 G. W. H. O. 2019. (World Health Organization, Geneva, 2019).

2 A. Uwimana et al. Emergence and clonal expansion of in vitro artemisinin-resistant Plasmodium falciparum kelch13 R561H mutant parasites in Rwanda. Nat Med, doi:10.1038/s41591-020-1005-2 (2020).

3 F. A. Siddiqui et al. Role of Plasmodium falciparum Kelch 13 Protein Mutations in P. falciparum Populations from Northeastern Myanmar in Mediating Artemisinin Resistance. mBio 11, doi:10.1128/mBio.01134-19 (2020).

4 J. A. Boddey &A. F. Cowman. Plasmodium Nesting: Remaking the Erythrocyte from the Inside Out. Annu Rev Microbiol 67, 243–269, doi:10.1146/annurev-micro-092412-155730 (2013).

5 N. J. Spillman, J. R. Beck &D. E. Goldberg. Protein Export into Malaria Parasite-Infected Erythrocytes: Mechanisms and Functional Consequences. Annu Rev Biochem 84, 813–841, doi:10.1146/annurev-biochem-060614-034157 (2015).

6 A. Alkhalil et al. Plasmodium falciparum likely encodes the principal anion channel on infected human erythrocytes. Blood 104, 4279–4286, doi:10.1182/blood-2004-05-2047 (2004).

7 S. A. Desai, S. M. Bezrukov &J. Zimmerberg. A voltage-dependent channel involved in nutrient uptake by red blood cells infected with the malaria parasite. Nature 406, 1001–1005, doi:10.1038/35023000 (2000).

8 A. Gupta et al. Complex nutrient channel phenotypes despite Mendelian inheritance in a Plasmodium falciparum genetic cross. PLoS Pathog 16, e1008363, doi:10.1371/journal.ppat.1008363 (2020).

9 D. Ito, M. A. Schureck &S. A. Desai. An essential dual-function complex mediates erythrocyte invasion and channel-mediated nutrient uptake in malaria parasites. Elife 6, doi:10.7554/eLife.23485 (2017).

10 W. Nguitragool et al. Malaria parasite clag3 genes determine channel-mediated nutrient uptake by infected red blood cells. Cell 145, 665–677, doi:10.1016/j.cell.2011.05.002 (2011).

11 P. Sharma, K. Rayavara, D. Ito, K. Basore &S. A. Desai. A CLAG3 mutation in an amphipathic transmembrane domain alters malaria parasite nutrient channels and confers leupeptin resistance. Infect Immun 83, 2566–2574, doi:10.1128/IAI.02966-14 (2015).

12 E. S. Sherling et al. The Plasmodium falciparum rhoptry protein RhopH3 plays essential roles in host cell invasion and nutrient uptake. Elife 6, doi:10.7554/eLife.23239 (2017).

13 J. A. Cooper et al. The 140/130/105 kilodalton protein complex in the rhoptries of Plasmodium falciparum consists of discrete polypeptides. Mol Biochem Parasitol 29, 251–260, doi:10.1016/0166-6851(88)90080-1 (1988).

14 H. J. Brown &R. L. Coppel. Primary structure of a Plasmodium falciparum rhoptry antigen. Mol Biochem Parasitol 49, 99–110, doi:10.1016/0166-6851(91)90133-q (1991).

15 O. Kaneko et al. The high molecular mass rhoptry protein, RhopH1, is encoded by members of the clag multigene family in Plasmodium falciparum and Plasmodium yoelii. Mol Biochem Parasitol 118, 223–231, doi:10.1016/s0166-6851(01)00391-7 (2001).

16 T. Y. Sam-Yellowe, H. Shio &M. E. Perkins. Secretion of Plasmodium falciparum rhoptry protein into the plasma membrane of host erythrocytes. J Cell Biol 106, 1507–1513, doi:10.1083/jcb.106.5.1507 (1988).

17 I. T. Ling et al. Characterisation of the rhoph2 gene of Plasmodium falciparum and Plasmodium yoelii. Mol Biochem Parasitol 127, 47–57, doi:10.1016/s0166-6851(02)00302-x (2003).

18 N. A. Counihan et al. Plasmodium falciparum parasites deploy RhopH2 into the host erythrocyte to obtain nutrients, grow and replicate. Elife 6, doi:10.7554/eLife.23217 (2017).

19 L. Vincensini, G. Fall, L. Berry, T. Blisnick &C. Braun Breton. The RhopH complex is transferred to the host cell cytoplasm following red blood cell invasion by Plasmodium falciparum. Mol Biochem Parasitol 160, 81–89, doi:10.1016/j.molbiopara.2008.04.002 (2008).

20 E. S. Sherling &C. van Ooij. Host cell remodeling by pathogens: the exomembrane system in Plasmodium-infected erythrocytes. FEMS Microbiol Rev 40, 701–721, doi:10.1093/femsre/fuw016 (2016).

21 C. M. Ho et al. Bottom-up structural proteomics: cryoEM of protein complexes enriched from the cellular milieu. Nat Methods 17, 79–85, doi:10.1038/s41592-019-0637-y (2020).

22 S. H. W. Scheres. RELION: Implementation of a Bayesian approach to cryo-EM structure determination. J Struct Biol 180, 519–530, doi:10.1016/j.jsb.2012.09.006 (2012).

23 A. Punjani, J. L. Rubinstein, D. J. Fleet &M. A. Brubaker. cryoSPARC: algorithms for rapid unsupervised cryo-EM structure determination. Nature Methods 14, 290-+, doi:10.1038/Nmeth.4169 (2017).

24 L. Swint-Kruse &C. S. Brown. Resmap: automated representation of macromolecular interfaces as two-dimensional networks. Bioinformatics 21, 3327–3328, doi:10.1093/bioinformatics.bti511 (2005).

25 E. Krissinel &K. Henrick. Inference of macromolecular assemblies from crystalline state. J Mol Biol 372, 774–797, doi:10.1016/j.jmb.2007.05.022 (2007).

26 A. Gupta et al. CLAG3 Self-Associates in Malaria Parasites and Quantitatively Determines Nutrient Uptake Channels at the Host Membrane. mBio 9, doi:10.1128/mBio.02293-17 (2018).

27 M. Dal Peraro &F. G. van der Goot. Pore-forming toxins: ancient, but never really out of fashion. Nat Rev Microbiol 14, 77–92, doi:10.1038/nrmicro.2015.3 (2016).

28 S. Mira-Martinez et al. Identification of Antimalarial Compounds That Require CLAG3 for Their Uptake by Plasmodium falciparum-Infected Erythrocytes. Antimicrob Agents Chemother 63, doi:10.1128/AAC.00052-19 (2019).

29 C. M. Ho et al. Malaria parasite translocon structure and mechanism of effector export. Nature 561, 70-+, doi:10.1038/s41586-018-0469-4 (2018).

30 J. W. H. M. Berman, Z. Feng, G. Gilliland, T.N. Bhat, H. Weissig, I.N. Shindyalov, P.E. Bourne. The Protein Data Bank. 28, 235–242 (2000).

31 J. R. Beck, V. Muralidharan, A. Oksman &D. E. Goldberg. PTEX component HSP101 mediates export of diverse malaria effectors into host erythrocytes. Nature 511, 592–595, doi:10.1038/nature13574 (2014).

32 M. E. Gonzalez &L. Carrasco. Viroporins. FEBS Lett 552, 28–34, doi:10.1016/s0014-5793(03)00780-4 (2003).

33 B. Martinac, Y. Saimi &C. Kung. Ion channels in microbes. Physiol Rev 88, 1449–1490, doi:10.1152/physrev.00005.2008 (2008).

34 N. J. Hardenbrook et al. Atomic structures of anthrax toxin protective antigen channels bound to partially unfolded lethal and edema factors. Nat Commun 11, 840, doi:10.1038/s41467-020-14658-6 (2020).

35 M. F. Horta. Pore-forming proteins in pathogenic protozoan parasites. Trends Microbiol 5, 363–366, doi:10.1016/S0966-842X(97)01109-8 (1997).

36 J. Chiu &P. J. Hogg. Allosteric disulfides: Sophisticated molecular structures enabling flexible protein regulation. J Biol Chem 294, 2949–2960, doi:10.1074/jbc.REV118.005604 (2019).

37 B. Schmidt, L. Ho &P. J. Hogg. Allosteric disulfide bonds. Biochemistry-Us 45, 7429–7433, doi:10.1021/bi0603064 (2006).

38 V. M. Kallakunta, A. Slama-Schwok &B. Mutus. Protein disulfide isomerase may facilitate the efflux of nitrite derived S-nitrosothiols from red blood cells. Redox Biol 1, 373–380, doi:10.1016/j.redox.2013.07.002 (2013).

39 G. N. Prado, J. R. Romero &A. Rivera. Endothelin-1 receptor antagonists regulate cell surface-associated protein disulfide isomerase in sickle cell disease. FASEB J 27, 4619–4629, doi:10.1096/fj.13-228577 (2013).

40 S. H. W. Scheres. A Bayesian View on Cryo-EM Structure Determination. J Mol Biol 415, 406–418, doi:10.1016/j.jmb.2011.11.010 (2012).

41 S. Q. Zheng et al. MotionCor2: anisotropic correction of beam-induced motion for improved cryo-electron microscopy. Nature Methods 14, 331–332, doi:10.1038/nmeth.4193 (2017).

42 A. Rohou &N. Grigorieff. CTFFIND4: Fast and accurate defocus estimation from electron micrographs. J Struct Biol 192, 216–221, doi:10.1016/j.jsb.2015.08.008 (2015).

43 K. Zhang. Gautomatch: a GPU-accelerated program for accurate, fast, flexible and fully automatic particle picking from cryo-EM micrographs with or without templates (2016).

44 E. F. Pettersen et al. UCSF chimera - A visualization system for exploratory research and analysis. J Comput Chem 25, 1605–1612, doi:10.1002/jcc.20084 (2004).

45 P. Emsley, B. Lohkamp, W. G. Scott &K. Cowtan. Features and development of Coot. Acta Crystallogr D 66, 486–501, doi:10.1107/S0907444910007493 (2010).

46 N. R. Coordinators. Database resources of the National Center for Biotechnology Information. Nucleic Acids Res 44, D7–D19, doi:10.1093/nar/gkv1290 (2016).

47 C. Aurrecoechea et al. PlasmoDB: a functional genomic database for malaria parasites. Nucleic Acids Res 37, D539–D543, doi:10.1093/nar/gkn814 (2009).

48 L. A. Kelley, S. Mezulis, C. M. Yates, M. N. Wass &M. J. E. Sternberg. The Phyre2 web portal for protein modeling, prediction and analysis. Nat Protoc 10, 845–858, doi:10.1038/nprot.2015.053 (2015).

49 P. D. Adams et al. PHENIX: a comprehensive Python-based system for macromolecular structure solution. Acta Crystallogr D 66, 213–221, doi:10.1107/S0907444909052925 (2010).

50 Schrodinger, LLC. The PyMOL Molecular Graphics System, Version 1.8 (2015).

51 A. Kucukelbir, F. J. Sigworth &H. D. Tagare. Quantifying the local resolution of cryo-EMEM density maps. Nature Methods 11, 63-+, doi:10.1038/Nmeth.2727 (2014).

52 V. B. Chen et al. MolProbity: all-atom structure validation for macromolecular crystallography. Acta Crystallogr D 66, 12–21, doi:10.1107/S0907444909042073 (2010).

